# Reprogramming Intrahepatic Cholangiocarcinoma Immune Microenvironment by Chemotherapy and CTLA-4 Blockade Enhances Anti-PD1 Therapy

**DOI:** 10.1101/2023.01.26.525680

**Authors:** Jiang Chen, Zohreh Amoozgar, Xin Liu, Shuichi Aoki, Zelong Liu, Sarah Shin, Aya Matsui, Zhangya Pu, Pin-Ji Lei, Meenal Datta, Lingling Zhu, Zhiping Ruan, Lei Shi, Daniel Staiculescu, Koetsu Inoue, Lance L. Munn, Dai Fukumura, Peigen Huang, Nabeel Bardeesy, Won Jin Ho, Rakesh. K. Jain, Dan G. Duda

## Abstract

Intrahepatic cholangiocarcinoma (ICC) has limited therapeutic options and a dismal prognosis. Anti-PD-L1 immunotherapy combined with gemcitabine/cisplatin chemotherapy has recently shown efficacy in biliary tract cancers, but responses are seen only in a minority of patients. Here, we studied the roles of anti-PD1 and anti-CTLA-4 immune checkpoint blockade (ICB) therapies when combined with gemcitabine/cisplatin and the mechanisms of treatment benefit in orthotopic murine ICC models. We evaluated the effects of the combined treatments on ICC vasculature and immune microenvironment using flow cytometry analysis, immunofluorescence, imaging mass cytometry, RNA-sequencing, qPCR, and *in vivo* T-cell depletion and CD8^+^ T-cell transfer using orthotopic ICC models and transgenic mice. Combining gemcitabine/cisplatin with anti-PD1 and anti-CTLA-4 antibodies led to substantial survival benefits and reduction of morbidity in two aggressive ICC models, which were ICB-resistant. Gemcitabine/cisplatin treatment increased the frequency of tumor-infiltrating lymphocytes and normalized the ICC vessels, and when combined with dual CTLA-4/PD1 blockade, increased the number of activated CD8^+^Cxcr3^+^IFN-γ^+^ T-cells. Depletion of CD8^+^ but not CD4^+^ T-cells compromised efficacy. Conversely, CD8^+^ T-cell transfer from *Cxcr3*^−/−^ versus *Cxcr3*^+/+^ mice into *Rag1*^−/−^ immunodeficient mice restored the anti-tumor effect of gemcitabine/cisplatin/ICB combination therapy. Finally, rational scheduling of the ICBs (anti-CTLA-4 “priming”) with chemotherapy and anti-PD1 therapy achieved equivalent efficacy with continuous dosing while reducing overall drug exposure. In summary, gemcitabine/cisplatin chemotherapy normalizes vessel structure, increases activated T-cell infiltration, and enhances anti-PD1/CTLA-4 immunotherapy efficacy in aggressive murine ICC. This combination approach should be clinically tested to overcome resistance to current therapies in ICC patients.

**One Sentence Summary:** Immune microenvironment reprogramming by chemotherapy and priming using CTLA-4 blockade render ICCs responsive to anti-PD-1 immunotherapy.

## INTRODUCTION

Intrahepatic cholangiocarcinoma (ICC) is an aggressive biliary tract cancer (BTC) characterized by late clinical presentation, frequent recurrence after local therapies, and resistance to systemic therapy (*1, 2*). The incidence of ICC has been increasing in the United States during the last two decades (*1, 3*). ICC arises in the liver, and its oncogenic drivers and microenvironment differ from extrahepatic biliary cancers, which is relevant for developing new therapeutic approaches. For patients diagnosed with BTC at advanced stages, systemic chemotherapy with gemcitabine and cisplatin (GC) has been the standard care over the last decade (*4–6*). This combination therapy provides a short delay in progression, but new therapeutic strategies are urgently needed because of the rapid development of resistance. Molecularly targeted therapies have been approved for specific ICC subsets recurring after GC, all of which have limited efficacy (*4, 5*). Overall, the 5-year survival for patients with ICC is a dismal 8%. Therefore, developing novel therapeutic strategies to improve ICC treatment efficacy remains a significant unmet need.

An attractive approach to impact ICC more broadly is using immune checkpoint blockade (ICB) therapy to reactivate and enhance anti-tumor immunity. The ability of ICBs—such as antibodies against programmed cell death protein (PD)-1 or its ligand PD-L1, or cytotoxic T lymphocyte antigen (CTLA)-4—to induce durable responses in advanced disease has established immunotherapy as a new pillar of cancer treatment (*7*). In BTC, a clinical trial of pembrolizumab—an anti-PD1 antibody—showed an overall response rate (ORR) of 17% (*8, 9*).Notably, a recent randomized phase III trial (TOPAZ-1) showed that GC chemotherapy with the PD-L1 antibody durvalumab demonstrated a hazard ratio of 0.80 for overall survival (OS) and an ORR of 27% versus 19% with GC alone as a first-line treatment for advanced BTC (*10*). Response rates increased to 27% from 19% of the patients. Interestingly, the addition of the CTLA-4 antibody tremelimumab to GC/durvalumab showed a 70% response rate in a phase II study in BTC patients (*11*).

These breakthroughs raise critical new questions about the mechanisms of the interaction between GC and ICB and their impact on the immunosuppressive microenvironment of ICC, a key mediator of treatment resistance. Previous studies demonstrated that chemotherapeutic drugs affect the viability of cancer cells and can also exert immunostimulatory effects. Some of these effects may be mediated by targeting immunosuppressive cells, while others may be mediated by increased immunogenicity secondary to cancer cell killing. However, a significant unmet need in this area of research has been the availability of animal models that reproduce the hallmarks of human ICC oncogenesis and microenvironment. It was reported that gemcitabine could deplete myeloid-derived suppressor cells (MDSCs) in several tumor-bearing animals and enhance antitumor immune activity (*12*). Gemcitabine can also polarize tumor-associated macrophages (TAM) towards antitumor phenotypes in pancreatic cancer (*13*). In addition to these direct immunostimulatory effects, gemcitabine and cisplatin have been reported to enhance the antigenicity and immunogenicity of different tumors by upregulating the expression of HLA class I in cancer cells (*14, 15*). Moreover, the expression of PD-L1 in cancer cells can be upregulated by cisplatin in human carcinomas (*16*). However, whether and how GC chemotherapy affects antitumor immunity in ICC remains unclear.

Previous studies demonstrated that, although PD-L1 is expressed in ICC cells, PD1-expressing lymphocytes infiltrate only the fibrous septa but not in tumor lobules (*17*). Findings corroborated these observations: T regulatory cells (Tregs) often accumulated in ICCs, most effector CD8^+^ CTLs and T helper cells were sequestered at the tumor margins (*18*). These features may explain the limited efficacy of using ICB alone in clinical trials. In addition, ICCs are poorly perfused and hypoxic, which contributes to their “immunologically cold” tumor microenvironments (*19–21*). These factors likely mediate resistance to ICB alone, as seen in most ICC patients (*22–24*). These limitations have shifted the focus on revealing and targeting the critical mechanisms of cancer immune evasion that are barriers to infiltration and activation of CTLs and ICB response. The infiltration of CD8^+^ CTLs is essential for effective cancer immunotherapy (*25, 26*). Chemokine receptor CXCR3 is expressed primarily on activated CD8^+^CTLs that produce perforin, granzyme, and interferon-γ (IFN-γ) (*27, 28*). The interaction of CXCR3 and its ligands, including interferon-inducible chemokines CXCL9, CXCL10, and CXCL11, is essential for the function of T cells. Although CXCR3 is not required for CD8^+^ T cell migration, it mediates intratumoral CD8^+^ T cell responses to anti-PD-1 therapy (*28*) and chemotherapy (*29*).

In the current study, we used two orthotopic ICC models that show characteristic oncogenic mutations, aggressive progression in the liver and at metastatic sites (lymph node and lung), development of ascites and pleural effusions, desmoplastic tumors with hypo-perfused vasculature and hypoxic microenvironment; and ICB resistance. We generated orthotopic tumors using cells established from two distinct genetically engineered mouse models: SS49 cells from *Idh2^R172K^/Kras^G12D^* mice and 425 cells from *TP53^KO^Kras^G12D^* mice (*30, 31*). Using these models, we evaluated the outcome of GC chemotherapy with single or dual ICB therapy in immunotherapy-resistant ICC models and their impact on the tumor microenvironment and anti-tumor immunity, as well as the potential mechanism of benefit.

## RESULTS

### Chemotherapy converts ICB-resistant ICCs to ICB-responsive tumors, and combination therapy increases survival

We first evaluated the *in vivo* efficacy of dual ICB (anti-PD1 and CTLA-4 antibodies) combined with GC chemotherapy versus each intervention type alone using *p53*-null (425) and *Idh*-mutant (SS49) murine ICC models (**Fig. 1a,b**). Mice received up to 6 injections of ICB and 9 injections of GC.

**Fig. 1:**
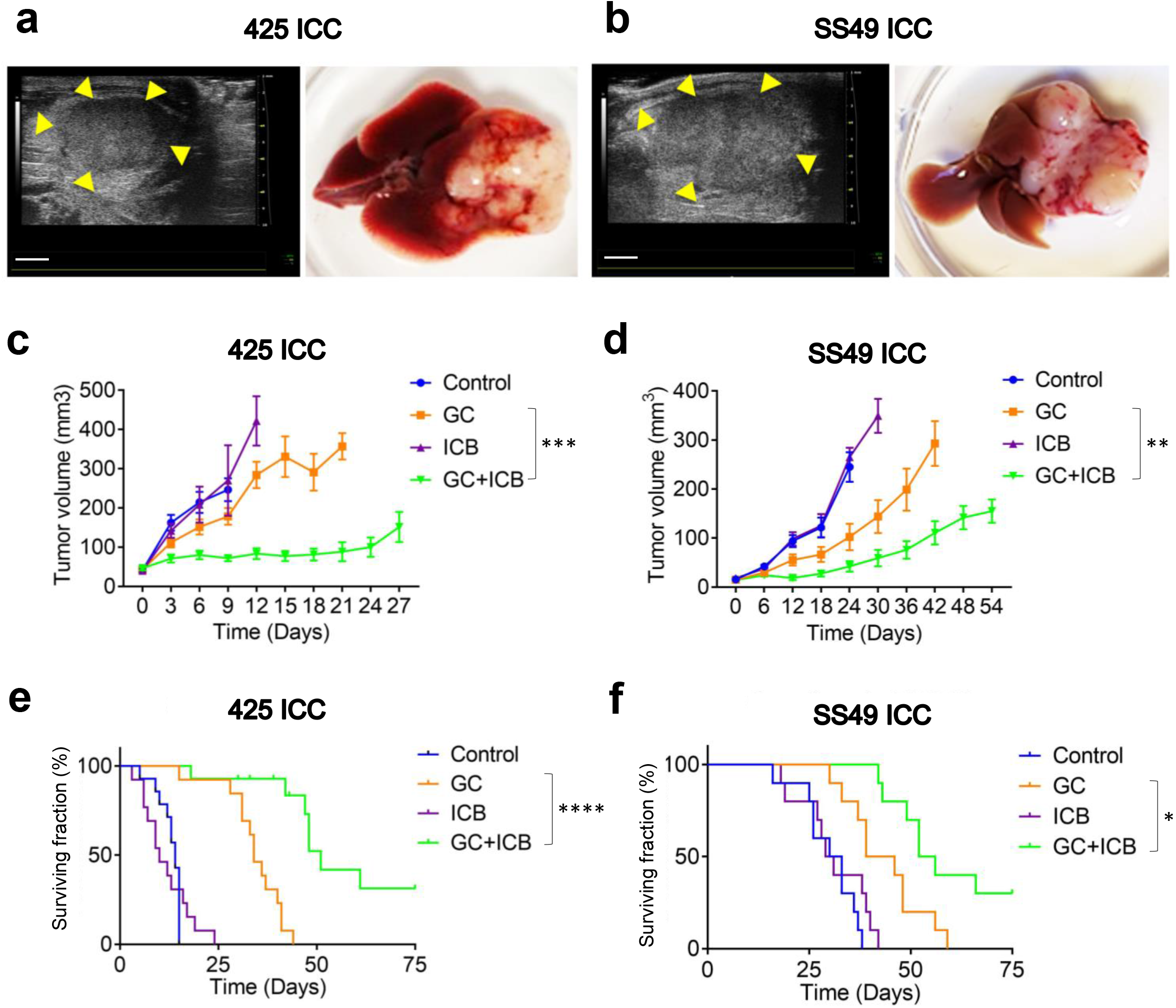
Standard chemotherapy converts ICB-resistant ICCs to ICB-responsive tumors, significantly delays tumor progression, and increases survival in mice. (**a**, **b**) Orthotopic ICC models in mice: *p53*^KO^*Kras*^G12D^ 425 murine ICC in C57Bl/6/FVB F1 mixed background mice (**a**) and *Idh2*^R172K^/*Kras*^G12D^ SS49 murine ICC in C57Bl/6 mice (**b**). Left, high-frequency ultrasound images. B-mode image of a Tumor 6 days after orthotopic implantation in a C57Bl/6/FVB F1 mouse and C57Bl/6 mouse. Scale bar: 2mm. Right, macroscopic appearance. (**c**, **d**) Tumor growth curves after treatment: GC+ICB therapy induced a tumor growth delay significantly superior to GC alone, while ICB alone was ineffective in 425 (**c**) and SS49 (**d**) murine ICC models. (**e**, **f**) Overall survival of ICC-bearing mice after treatment: GC+ICB therapy induced a survival advantage that was significantly superior to GC alone, while ICB alone showed no efficacy in the 425 and SS49 (**f**) murine ICC models. GC, gemcitabine plus cisplatin; ICB, immune checkpoint blockade (anti-PD1 antibody plus anti-CTLA-4 antibody). *p<0.05; **p<0.01; ***p<0.001; ****p<0.0001 from Dunn’tt’s multiple comparisons tests (**c**, **d**) and Log-rank (Mantel-Cox) test (**e**, **f**). *In vivo* studies were performed in duplicate; n=14 mice (425 murine ICC) and n=10 (SS49 murine ICC).

We found that all combination treatments were feasible in these models. These tumors were resistant to dual ICB therapy alone; however, combining GC with dual ICB significantly delayed the growth of established ICCs in both models (**Fig. 1c,d** and **Fig. S1a,b**). Moreover, the combination of GC and dual ICB substantially improved median OS compared to chemotherapy alone in mice bearing these aggressive ICCs (**Fig. 1e,f**). In addition, combined GC/dual ICB treatment significantly reduced the formation of lung metastases and disease-related morbidity, i.e., the incidence of bloody ascites and pleural effusions (**Fig. S1c-h**). Combination therapy was associated with weight loss in some mice but less than the 15% specified per protocol (**Fig. S1i,j**). These data showed that standard GC chemotherapy could convert *p53*-null and *Idh*-mutant ICCs to dual ICB therapy-responsive tumors and that combination therapy is feasible.

### GC/ICB therapy reprograms the immune microenvironment of the ICC

In separate experiments, we examined the effect of GC/ICB combination therapy on the immune microenvironment of ICC. To this end, we conducted a time-matched study in ICC-bearing mice. All mice were sacrificed after 8 days of treatment with dual ICB (anti-PD1 and CTLA-4 antibodies) combined with GC chemotherapy when tumor growth was already significantly delayed compared to each intervention alone. Immune cell infiltration was measured by immunofluorescence (IF) in tumor sections and flow cytometry in dissociated tumor tissues. GC chemotherapy treatment increased CD8^+^ T-cell infiltration and proliferation/activation in ICC tissue (**Fig. 2a,c**). Since these murine ICCs are typically hypo-perfused due to blood vessel abnormalities (*32*), we also used IF for CD31 (an endothelial marker) and α-SMA^+^ (a perivascular cell marker) to evaluate changes in vascular density. We found that GC increased coverage by perivascular cells, which conferred vessel stability and maturity – hallmarks of vascular normalization (**Fig. 2b,d,e**).

**Figure 2:**
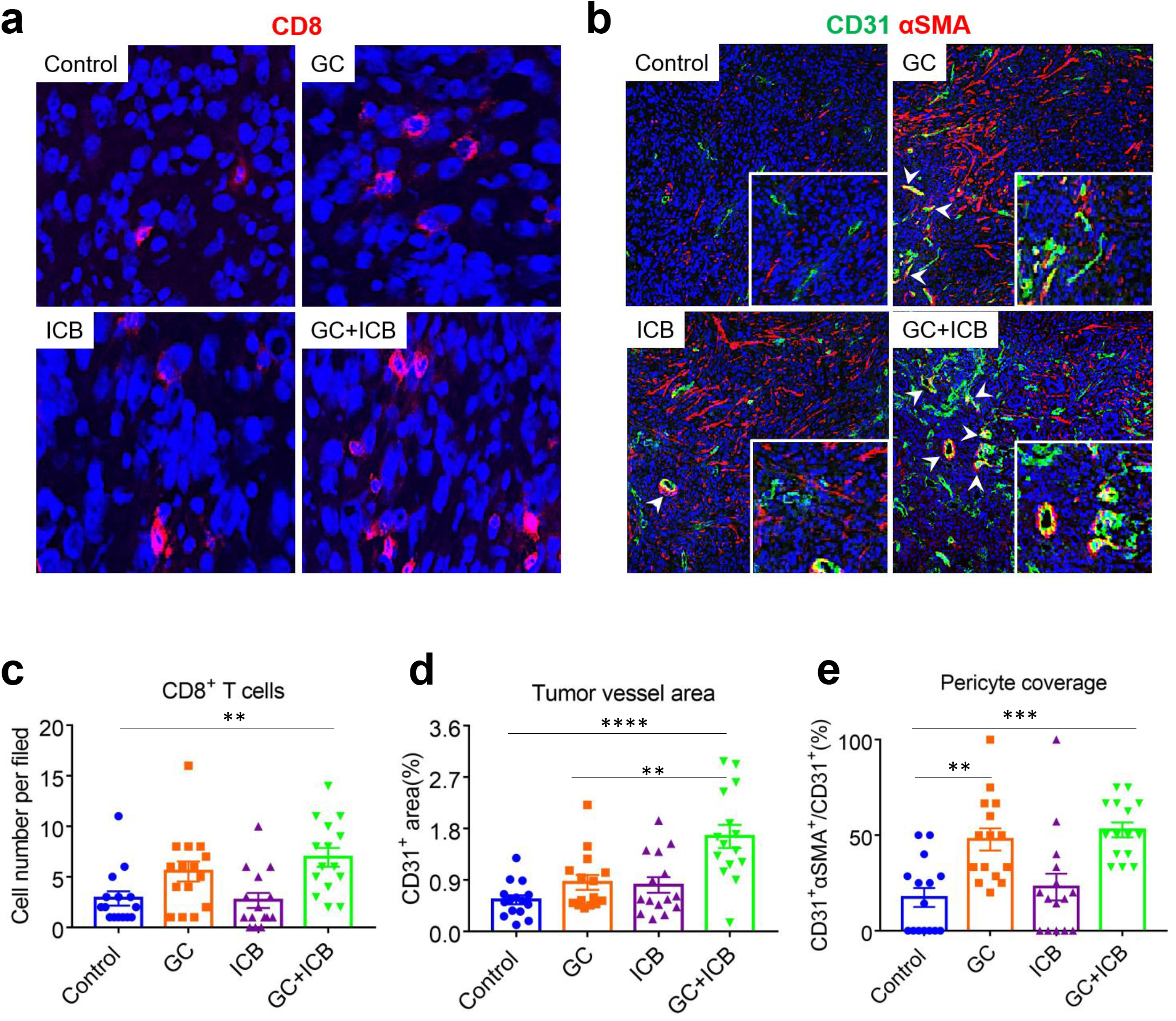
GC chemotherapy increases the infiltration of CD8^+^ T and normalizes tumor vasculature in 425 murine ICC model. (**a**, **b**) Representative immunofluorescence (IF) for the T-cell marker CD8 (**a**) and the endothelial and perivascular cell markers CD31 and α-SMA, respectively (**b**). (**c**) ICC tissue infiltration by CD8^+^ T-cells was increased in mice treated with GC alone or with ICB. (**d**, **e**) GC can increase CD31^+^ in the tumor and induce normalization of ICC vessels, both of which were quantified by immunofluorescence in ICC tissue. GC: gemcitabine plus cisplatin; ICB: anti-PD1 antibody plus anti-CTLA-4 antibody. **p<0.01; ***p<0.001; ****p<0.0001 from Tukey’s multiple comparisons test. In vivo studies were performed in duplicate; n=15 mice per group.

Of note, prior studies demonstrated that vascular normalization increases the recruitment of effector T-cells and the efficacy of immunotherapy (*33*). T-cells can also normalize tumor vessels and mediate anti-tumor immunity (*34*). GC/dual ICB therapy increased ICC’s CTL infiltration, proliferation, and normalized vessel structure.

### CTLA-4 blockade mediates the efficacy of combined ICB/GC therapy in ICB-resistant 425 ICC model

Next, we tested the specific roles of CTLA-4 and PD1 blockade in the efficacy of combined GC/ICB therapy in mice with established orthotopic murine 425 ICC. To this end, we evaluated the *in vivo* efficacy of treatment with: (**i**) GC and dual ICB (anti-PD1 and anti-CTLA-4 antibodies), (**ii**) GC with anti-PD1 antibody, (**iii**) GC with anti-CTLA-4 antibody, (**iv**) GC chemotherapy alone, (**v**) dual ICB alone, (**vi**) anti-PD1 antibody alone, (**vii**) anti-CTLA-4 antibody alone, or (**viii**) isotype-matched IgG control. All agents were administered until mice became moribund, tumors reached the maximum size allowed per protocol, or tumors became undetectable by imaging. We found that GC treatment combined with anti-PD1 therapy induced a growth delay and survival advantage that was not superior to GC alone in this model. In contrast, the combination of GC and anti-CTLA-4 treatment caused a significant delay in tumor growth and an increase in median OS compared to GC alone treatment, which was inferior to GC with dual ICB in this ICC model (**Fig. 3a,b** and **Fig. S2a**).

**Figure 3:**
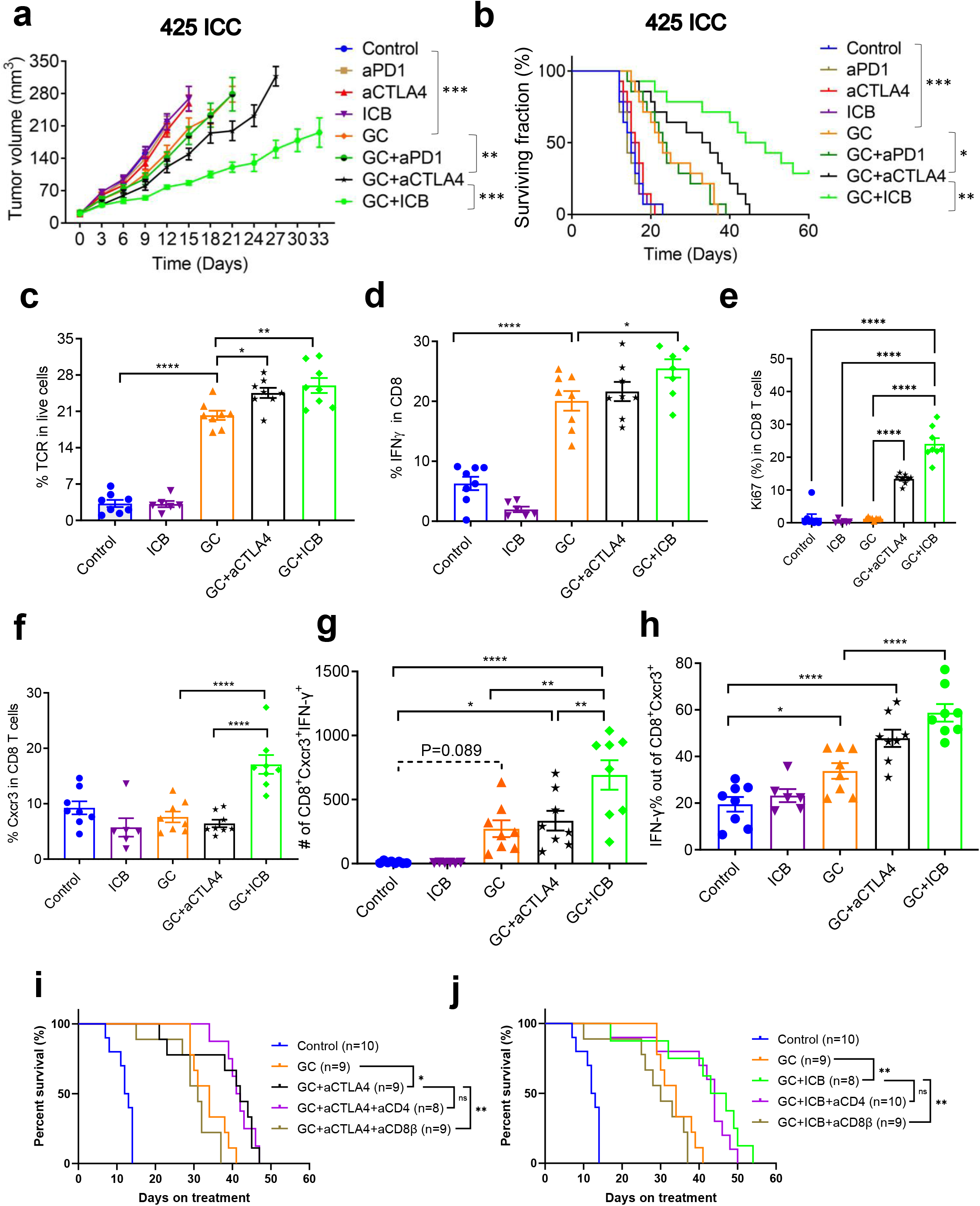
CTLA-4 blockade mediates the efficacy of GC/ICB therapy in ICC and increases CD8^+^Cxcr3^+^ CTL frequency in murine ICC. (**a**, **b**) Tumor growth kinetics and survival distributions after treatment in orthotopic 425 murine ICC model: GC/anti-CTLA4 Ab combination is significantly superior to GC alone and in delaying tumor growth (**a**) and increasing survival (**b**); addition of anti-PD1 Ab to GC/anti-CTLA4 Ab (GC/ICB) but not to GC alone induces a more significant delay in tumor growth (**a**) and increase in survival (**b**). (**c-h**) Immunophenotyping of treated ICC tissues at days 10 (control and ICB groups) and 20 (GC-containing groups): TCR^+^ tumor-infiltrating lymphocyte (TIL) frequencies were higher in all GC-treated groups and significantly increased in the groups receiving GC and anti-CTLA4 Ab (**c**). CD8^+^IFN-γ^+^ T-cell frequencies were higher in all GC-treated groups and significantly increased in the GC/ICB group (p=0.0044 vs. GC alone) (**d**). Ki67^+^ CD8^+^ T-cell frequency (**e**), CD8^+^Cxcr3^+^ T-cell frequencies (**f**), CD8^+^Cxcr3^+^IFN-γ^+^ T cell numbers (**g**), and frequencies (**h**) were significantly increased in the GC/ICB group (p=0.0002 vs. GC alone, p=0.0217 vs. GC+aCTLA4). (**i,j**) CD8^+^ T cells depletion rather than CD4^+^ T cells depletion compromised the survival benefit in both GC/anti-CTLA4 and GC/dual ICB treatment. *p<0.05; **p<0.01; ****p<0.0001 from Dunnett’s multiple comparisons tests (**a**) and Log-rank (Mantel-Cox) test (**b, i,j**) and Tukey’s multiple comparisons tests (**c**-**h**). *In vivo* studies were performed in duplicate; n=14 mice per group.

Prolonged, continuous administration of combination therapy induced toxicity in this experiment, including weight loss leading to treatment breaks in some mice (**Fig. S2b**). The effects of treatment on the inhibition of lung metastases were significant only after combination therapy of GC with anti-CTLA-4 or GC with dual ICB (**Fig. S2c**). Similarly, disease-related morbidity was reduced in all GC-treated groups, and to the greatest extent, in the GC combined with the dual ICB group (**Fig. S2d,e**). These data demonstrate that CTLA-4 blockade is essential for the efficacy of combination GC/dual ICB treatment in an ICC model, which is resistant to immunotherapy alone and can render these tumors responsive to anti-PD1 therapy.

### Combining GC chemotherapy with dual ICB treatment increases intratumoral CD8^+^ CTL infiltration and activation

Next, to decipher the mechanism of benefit of GC/ICB therapy, we first repeated the time-matched experiments to evaluate the effects of combination therapy by immune profiling of responding 425 murine ICCs at day 20, when the differences in growth delay were significant between groups (**Fig. S3a**). In this experiment, ICC-bearing C57Bl/6/FVB F1 mice were treated with: (**i**) GC alone, (**ii**) dual ICB alone, (**iii**) GC plus anti-CTLA-4 antibody, (**iv**) GC combined with dual ICB, or **v**) IgG control. Mice from the control group and dual ICB alone had to be sacrificed on day 10 due to rapid tumor progression; these tumor tissues were collected and used as a reference. Tissues were digested for flow cytometry and embedded in paraffin for imaging mass cytometry (IMC).

To evaluate the overall frequency of infiltrating T lymphocytes, we counted the T-cell receptor (TCR)^+^ cells by flow cytometry in enzymatically digested ICC tissues. We found significantly higher frequencies of tumor-infiltrating lymphocytes (TILs) in all GC-treated groups; moreover, there was a significant increase in the fraction of infiltrating lymphocytes after GC with CTLA-4 antibody and GC with dual ICB versus GC alone group (**Fig. 3c** and **Fig. S3a**). Among TILs, approximately 60% were CD8^+^ T-cells in all groups (**Fig. S3b,c**). Moreover, the fraction of CD8^+^interferon-gamma (IFN-γ)^+^ T-cells was higher in the tumors from GC-treated groups and was significantly increased after GC with dual ICB (p<0.05 versus GC alone) (**Fig. 3d** and **Fig. S3d**). Notably, the fraction and the number of Ki67^+^CD8^+^ T cells were significantly higher in tumors after GC/anti-CTLA4 and GC/ICB treatment (**Fig. 3e** and **Fig. S3e**). Ki67^+^CD4^+^conventional T cells also increased after GC/anti-CTLA4 and GC/dual ICB treatment. Of note, the frequency of CD4^+^ cells was substantially lower than that of CD8^+^ T cells (**Fig. S3f**).

Among all markers of T-cell activation tested (IL-2, IL-12, IFN-γ, Cxcr3), we found that the frequency and number of CD8^+^Cxcr3^+^ and CD8^+^Cxcr3^+^IFN-γ^+^ T-cells were significantly increased in tumors treated with GC combined with dual ICB compared to GC alone or GC combined with anti-CTLA-4 antibody (**Fig. 3f-h** and **Fig. S3g,h**). Of note, we also found significant increases in CD4^+^FoxP3^+^ Tregs, resulting in comparable CD8^+^ T-cell to Treg ratios between GC-treated groups, but the absolute counts of Tregs were low (**Fig. S3i-l**). Further phenotyping of CD8^+^ T cells showed that GC/ICB combination therapy reduced the markers of immunosuppression (markers associated with CD8^+^ T cells exhaustion), including PD1, TIGIT, Tim3, GITR, Vista, and Lag3 and increased markers that indicate high immunity such as CD44 marking effector function and CD69, an early activation marker (**Fig. S4**).

IMC analyses confirmed the significant increase in intratumoral CD8^+^ T cells and CD31^+^ vessel area in ICC tissues from the GC/dual ICB group (**Fig. S5a-f**). Moreover, CD3^+^TCF1^+^, as well as CD8^+^CD3^+^TCF1^+^ infiltration, was significantly increased in ICCs from the GC/dual ICB treatment group (**Fig. S5g,h**).

These results indicate that adding CTLA-4 blockade to GC increases the number of activated CTLs and that the addition of PD1 blockade further increases the fraction of CD8^+^Cxcr3^+^IFN-γ^+^activated T-cells in ICC tissue. They also raise the question of whether the expression of Cxcr3 ligands changes after GC/ICB treatment.

### Expression of Cxcr3 ligands is increased after combined GC and dual ICB treatment in ICC tissues

Using tissues collected in a time-matched manner, we employed RNA sequencing (RNA-Seq) to evaluate the transcriptional changes in murine 425 ICC tissues after 8 days of treatment with GC alone, dual ICB alone, and GC with dual ICB versus IgG control. Bulk RNAseq analysis showed significant increases in the expression of over 1,200 genes in the GC/ICB group compared to the other groups (**Fig. S6a** and **Dataset S1**). Gene enrichment analyses showed significant activation of pathways related to cellular immunity (**Fig. S6b**). Of these, the transcription of Cxcl9, Cxcl10, and Cxcl11, as well as their receptor Cxcr3, increased after combination therapy versus control (n=3) (**Fig. S7a**-**d**). The activation of this chemokine pathway is critical for CTL function and response to anti-PD1 therapy (*28, 35, 36*). We used qPCR analysis to validate the increase in Cxcl10 and Cxcl11 expression and found a trend for increased Cxcl9 and Cxcr3 expression at this early time-point (n=5) (**Fig. S7e-h**).

### Cxcr3^+^CD8^+^ T-cells mediate the benefit of GC/ICB combination therapy in murine ICC

Next, we performed depletion experiments to determine whether CD4^+^ or CD8^+^ T-cells mediate the benefit of combination therapy. CD4^+^ cell depletion did not affect median OS after GC/CTLA-4 Ab or GC/ICB therapy, whereas CD8^+^ cell depletion completely compromised their efficacy **(Fig. 3i,j**). Median OS was 42 days in the GC/CTLA-4 Ab group versus 31 days in the survival of GC/CTLA-4 Ab/aCD8β Ab group (p=0.0046) and 45 days in the GC/ICB group versus 30 days in the GC/ICB/aCD8β Ab group (p=0.0048).

Next, we used the adoptive T-cell transfer model to establish whether Cxcr3^+^CD8^+^ T-cells mediate the efficacy of GC/ICB therapy in the 425 murine ICC model. We first transferred one million CD8^+^ T-cells from *Cxcr3*^+/+^/C57Bl/6 or *Cxcr3*^−/−^/C57Bl/6 to *Rag1*^−/−^/C57Bl/6 mice, which lack functional T-cells. Two days after the T-cell transfer, we orthotopically implanted murine 425 ICC cells in the *Rag1*^−/−^/C57Bl/6 mice. When tumors were established, the mice received GC/dual ICB combination therapy or GC alone, and tumor growth was monitored by ultrasound imaging (**Fig. S8**).

We found that ICC growth after GC therapy was not different between mice receiving CD8^+^ T-cell transfer from *Cxcr3*^+/+^/C57Bl/6 versus *Cxcr3*^−/−^/C57BL/6 donor mice. In contrast, the efficacy of GC/ICB therapy was significantly diminished in mice that received CD8^+^ T-cells from *Cxcr3*^−/−^/C57Bl/6 versus those which received CD8^+^ T-cells from *Cxcr3*^+/+^/C57Bl/6 mice, with an inferior tumor growth delay and survival benefit (**Fig. 4a**,**b** and **Fig. S9a**). Treatment was not associated with significant weight loss in any groups (**Fig. S9b**). Moreover, the metastatic lung burden was lowest in mice treated with GC/ICB, which received CD8^+^ T-cell transfer from *Cxcr3*^+/+^/C57BL/6 mice (**Fig. S9c**). Finally, in contrast to the other treatment groups, these mice were also free of bloody ascites and pleural effusions (**Fig. S9d**,**e**).

**Figure 4:**
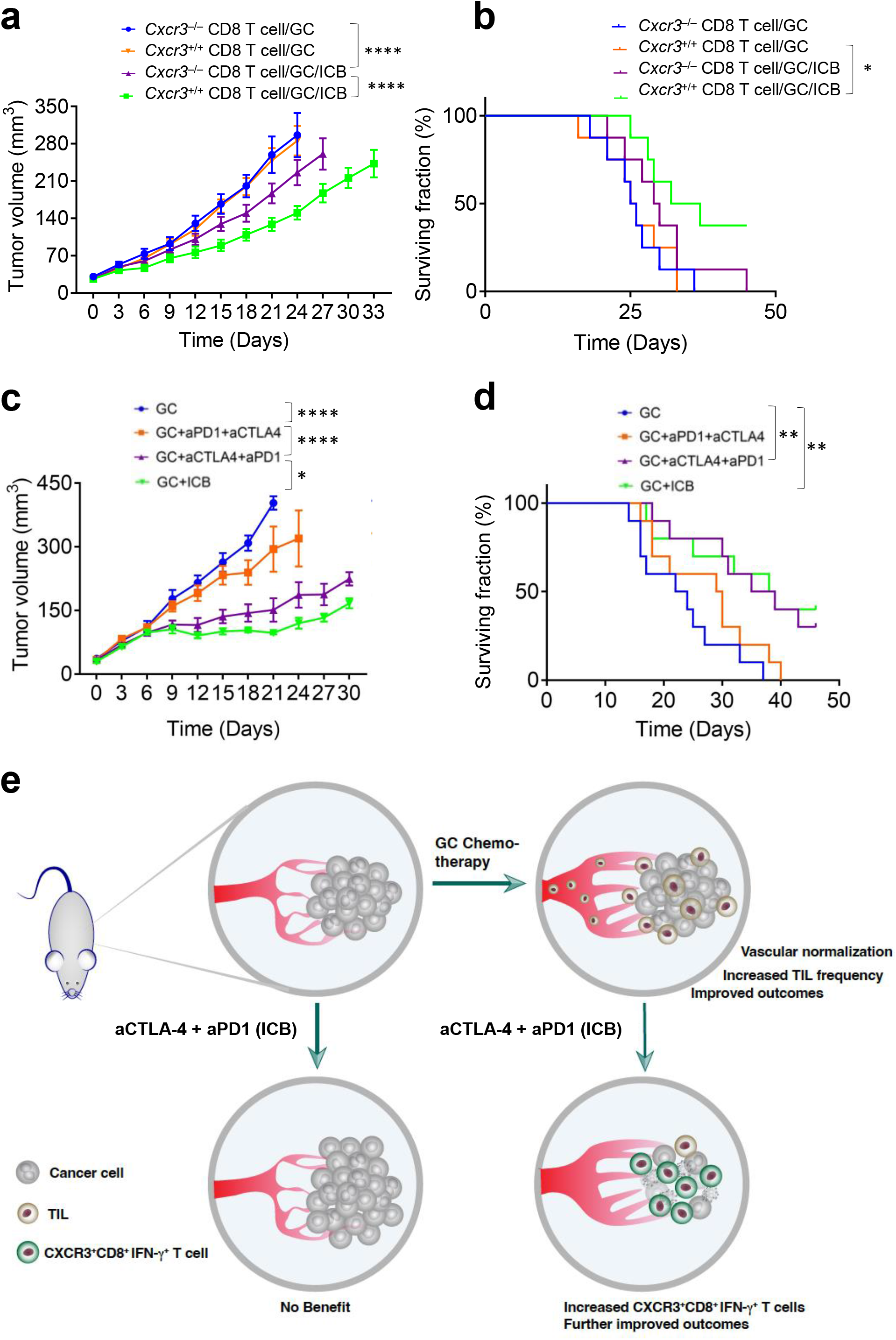
CD8^+^Cxcr3^+^ T cells and anti-CTLA-4 therapy priming mediate the benefit of GC/ICB combination therapy in 425 murine ICC. (**a, b**) The tumor growth delay (a) and overall survival (b) benefit of GC+ICB over GC alone therapy are significantly reduced in *Rag1*^−/−^/C57Bl/6 mice which received CD8^+^ T-cell transfer from *Cxcr3*^−/−^/C57Bl/6 mice transferred to mice bearing ICC (n=8 mice). (c, d) Tumor growth delay (c) and overall survival (d) after treatment with GC+anti-CTLA-4 priming for a week followed by GC+anti-PD1 maintenance were comparable to full dose GC/ICB and significantly superior to GC+anti-PD1 priming for a week followed by GC+anti-CTLA-4 maintenance and to GC alone (n=10). (e) Proposed mechanism of benefit: Standard gemcitabine/cisplatin therapy reprograms the immune microenvironment and normalizes vessels in ICB-resistant ICC, and combination with anti-PD1/CTLA-4 dual ICB therapy increased efficacy mediated by anti-CTLA-4 therapy priming and tumor infiltration by activated CD8^+^Cxcr3^+^ T-cells. * P<0.05; ** P<0.01; **** P<0.0001 from Dunnett’s multiple comparisons tests (a, c) and Log-rank (Mantel-Cox) test (b, d).

These data show that the benefits of GC combined with anti-CTLA-4 Ab alone or dual ICB partly depended on CD8^+^Cxcr3^+^ T-cells.

### ICB scheduling with chemotherapy can achieve efficacy while reducing drug exposure

The GC/ICB combination showed efficacy but was associated with adverse effects in continuous administration. Thus, we used the 425 murine ICC orthotopic model to test the effect of treatment de-escalation based on scheduling the agents to leverage their mechanisms of action on efficacy. To this end, mice with established tumors were randomized into one of the following treatment groups: (**i**) GC until endpoint (maximum chemo-drugs exposure); (**ii**) GC with anti-CTLA-4 and anti-PD1 antibodies until endpoint (maximum chemo- and ICB exposure); (**iii**) GC until endpoint with 1 week of anti-CTLA-4 Ab and maintenance anti-PD1 Ab from day 8 to endpoint (chemo/CTLA-4 blockade “priming”/reduced ICB drug exposure); and (**iv**) GC until endpoint with anti-PD1 antibody for the first week and anti-CTLA-4 Ab from day 8 to endpoint (chemo/PD1 blockade “priming”/reduced ICB drug exposure) (**Fig. S10a**). We found that the efficacy of ICB “priming” with anti-CTLA-4 antibody but not anti-PD1 antibody was equivalent to that of maximum chemotherapy and ICB drug exposure in terms of tumor growth delay and OS benefit (**Fig. 4c,d** and **Fig. S10b**). Moreover, anti-CTLA-4 antibody priming was associated with lower toxicities than in mice from the maximum drug exposure group while showing equally low ratios of lung metastasis, ascites, and pleural effusions (**Fig. S10c-f**).

## DISCUSSION

Over the last decade, ICBs have revolutionized systemic therapy for cancer after inducing durable responses in some malignancies. Unfortunately, these cases represent a minority (typically 20-30%) for most tumor types (*37*). Using orthotopic ICC models, we show that GC chemotherapy converts ICCs from immunologically “cold” tumors into “hot” tumors by increasing effector CD8^+^ T-cell recruitment facilitated by induction of vascular normalization. These effects enhanced the efficacy of immunotherapy with ICB via recruitment of activated CD8^+^Cxcr3^+^ T-cells, leading to increased survival and reduced morbidity in two aggressive orthotopic models of ICC. Our study reveals that standard chemotherapy for ICC facilitates anti-CTLA-4-based ICB therapies and the mechanisms of benefit for these treatment interactions (**Fig. 4e**).

Vascular normalization by GC chemotherapy in ICC may be indirectly mediated by killing cancer cells, the key source of pro-angiogenic factors. This concept was first demonstrated by hormone withdrawal in a hormone-dependent tumor model (*38*). GC treatment also increases the accumulation of CD8^+^ T cells and, when combined with dual CTLA-4 and PD1 blockade, showed durable responses mediated partly by activation of CD8^+^Cxcr3^+^ CTLs. IMC analyses confirmed the increased vascularization and T-cell infiltration, associated with a reduction in the number of cancer cells in the murine ICC after GC and dual ICB combination treatment, consistent with IF and flow cytometric studies.

CTLA-4 blockade played a critical role when combined with GC in tumor responses and induced an increase in activated CD8^+^ T cells despite the persistence of Tregs, which were comparatively less frequent. Given the recent data showing the feasibility and efficacy of GC chemotherapy with anti-PD-L1 therapy in BTC, the approach tested here is immediately applicable as combination therapy in ICC. The concept of using combination therapy to address resistance to anti-PD1/PD-L1 therapy is widely accepted and is actively pursued in clinical studies (*39–41*). However, its implementation is limited by an incomplete understanding of treatment interactions in specific tumor contexts, serious toxicity concerns, and high economic costs. Our study demonstrates the efficacy of mechanism-based multimodality therapy for ICC, which minimizes drug exposure for the reagents used in combined strategies while maintaining optimal effectiveness.

Although early studies of Cxcr3 focused on its role in driving the recruitment of activated T cells into inflamed tissues, Cxcr3 was identified as an activation marker on T cells (*28, 42–45*). For CD8^+^ T cells, Cxcr3 might be involved in the directed migration of these cells and their activation and proliferation in tumors (*46*). Our study demonstrates that the reprogramming of the ICC immune microenvironment by chemotherapy and the enhancement of CTLA-4 blockade of anti-PD1 treatment efficacy is in part dependent on Cxcr3^+^CD8^+^ T-cells. Our immune profiling results show that, while CTLA-4 blockade alone did not significantly enhance the T-cell infiltration induced by GC chemotherapy, it increased the fraction of proliferating CD8^+^ T-cells and treatment efficacy. Moreover, when added to GC/anti-PD1 therapy, CTLA-4 blockade significantly increased the infiltration and activation of CD8^+^ T cells. While ICC infiltration by CD4^+^ conventional T-cells also increased after GC/dual ICB treatment, a similar trend was seen for Tregs. Depletion experiments demonstrated that treatment efficacy depended on CD8^+^ and not CD4^+^ T-cells. These results are consistent with a comparatively lower frequency of CD4+ versus CD8+ T-cells in the ICC tissues.

Among the CD8^+^ T-cells subsets infiltrating ICC tissue after GC/dual ICB treatment, our IMC analyses showed an increase in CD8^+^TCF1^+^ and CD8^+^GZMB^+^ T-cells, known to be associated with the activation and recruitment of CD8^+^Cxcr3^+^ T cells (*45*). Moreover, we found an increased frequency of CD44^+^ and CD69^+^ CD8^+^ T-cells in the ICC microenvironment, consistent with early activation of effector T-cells. CD8^+^ T-cell expansion seen after the addition of CTLA-4 blockade alone to GC was also associated with the expression of cell exhaustion markers, including PD1, TIGIT, Tim3, and Lag3. Importantly, when GC was combined with dual CTLA-4/PD-1 blockade, expression of these immunosuppressive markers decreased on the infiltrating CD8^+^ T-cells. These results demonstrate the role of CTLA-4 blockade and the importance of PD1 blockade in efficacious ICB therapy with GC in ICC. Based on these data, we postulate that the rational scheduling of the ICBs (anti-CTLA-4 “priming”) with GC chemotherapy and anti-PD1 therapy can achieve equivalent efficacy with continuous dosing while reducing overall drug exposure in ICC patients.

In summary, we demonstrate that GC chemotherapy can normalize vessel structure and increase activated T-cell infiltration in ICC. GC combined with anti-PD1/CTLA-4 immunotherapy was feasible and showed efficacy in murine ICCs resistant to ICB or GC/anti-PD1 therapy. These new insights into chemotherapy/ICB treatment interactions may directly inform clinical trials designed to increase efficacy while reducing drug exposure in ICC patients.

## MATERIALS AND METHODS

### Cells and culture condition

We used the murine ICC cell lines SS49 and 425 cells established from spontaneous tumors in *Idh2^R172K^/Kras^G12D^* and *p53^−/−^Kras^G12D^* mice, respectively (*30, 31*). Cells were cultured in Dulbecco’s Modified Eagle’s Medium (DMEM) (ThermoFisher, USA) with 10% fetal bovine serum (FBS) (Hyclone, SH30071.03) and 10% penicillin-streptomycin in 5% CO2 at 37 °C. Mycoplasma contamination and cell authentication were routinely performed before *in vivo* studies.

### ICC mouse models

Orthotopic ICCs were induced by grafting cells in the liver of C57Bl/6/FVB F1 mixed background mice and *Rag1*^−/−^/C57Bl/6 mice using 425 cell line and C57Bl/6 wild-type for SS49 cells. The suspensions of 425 cells or SS49 cells and mixed with Matrigels (1:1 (volume/volume with the cell suspension)) were injected into the subcapsular region of the mesolimbic liver parenchyma using small 0.5ml syringes with 28.5 gauge needles. To avoid leakage of tumor cells from the injection sites, which might lead to local spread and ‘seeding’ of metastasis in the peritoneal cavity, we limited the injection volume to 20 μl (10^6^ cells in 20 μl per mouse). In addition, a steady and slow injection was performed to prevent leakage of the injected cell suspension and to minimize the damage to the surrounding liver tissues. After the removal of the needle, the liver surface at the site of the needle tract was covered with Gelfoam for 5 min to reduce bleeding and potential backflow (*31, 32*). All treatments were initiated in mice with established tumors when the ICCs reached 5mm in diameter, measured by high-frequency ultrasound imaging. Tumor growth and treatment response were also monitored by ultrasound imaging (**Fig. 1a,b**). The *p53*-null (425) and *Idh-*mutant (SS49) murine ICC models showed reproducible aggressive primary tumor growth and spontaneous lung metastases. Spontaneous metastatic burden was evaluated by considering the number and the size of the nodules, as previously described (*32*). For survival studies, moribund status was used as the endpoint per protocol, defined as symptoms of prolonged distress, >15% weight loss compared with the starting date, body condition score >2, and tumor size of >15mm in diameter. All experiments were performed under the MGH IACUC-approved protocol 2014N000083 titled “Targeted treatments for cholangiocarcinoma”.

### Imaging of orthotopic intrahepatic cholangiocarcinoma

Tumor growth and treatment response were monitored by high-frequency ultrasound imaging. For this model’s longitudinal evaluation of tumor growth, we used an ultrasound device equipped with specific probes for small-animal imaging at appropriate time points (6 days after implantation and then every 3 days (for the 425 model) and six days (for the SS49 model), was used to longitudinally assess tumor growth noninvasively under isoflurane anesthesia (*32*).

### Treatments

Gemcitabine and cisplatin were purchased from Pfizer and Fresenius Kabi, respectively. Mouse anti-PD1 antibody (clone RMP-014), anti-CTLA-4 (clone 9D9), anti-CD4 (clone GK1.5), and anti-CD8β (clone 53-5.8) were purchased from BioXcell (Lebanon, NH). Gemcitabine and cisplatin were administered intraperitoneally (i.p., 60 mg/kg for gemcitabine and 0.3 mg/kg for cisplatin twice/week), anti-mouse PD1 antibody and anti-CTLA-4 antibody were administered by retrobulbar injection (10 mg/kg, every three days), and anti-CD4 and anti-CD8β depleting antibodies were administered by intraperitoneal injection (10 mg/kg, thrice a week).

For adoptive cell transfer studies, we enriched CD8^+^ T-cells from *Cxcr3*^−/−^/C57Bl/6 mice or non-transgenic C57Bl/6 mice to 80% purity using a negative selection method (Stemcell Technologies, Vancouver, Canada). Then, cells were sorted to 98% purity using AriaII FACS sorter in RPMI/with 10% FBS. Cells were washed with PBS twice and concentrated to 1×10^6^ cells per 100μl. Cells were transferred via tail vein to *Rag1*^−/−^/C57Bl/6 mice two days before the ICC implantation into the liver.

### Immunofluorescence (IF)

Six-μm-thick frozen sections of ICC tissue were prepared for IF. We used an anti-CD31 antibody to identify endothelial cells, an anti-α-SMA antibody to identify perivascular cells, and an anti-CD8 antibody for staining T-cells. All primary antibodies are listed in **Table S1**. All secondary antibodies were purchased from Jackson ImmunoResearch (West Grove, PA, USA). Frozen sections from OCT-embedded tissue blocks were washed with PBS and treated with normal donkey serum for blocking. Primary antibodies were applied overnight at 4°C, followed by the reaction with appropriate secondary antibodies for 2hr at 24°C. Analysis was performed in five random fields in the tumor tissues under ×400 magnification using a laser-scanning confocal microscope (Olympus, FV-1000). Data were analyzed using ImageJ (US NIH) and Photoshop (Adobe Systems Inc.) software.

### Quantitative real-time reverse transcription-polymerase chain reaction (qPCR)

Total RNA was isolated using Rneasy Mini Kit (Qiagen Inc.) and analyzed by the NanoDrop system. qPCR was carried out using iTaq Universal SYBR Green Supermix (Bio-Rad, Inc.) and was amplified at the annealing temperature of 60°C with the primers (**Table S2**). GAPDH was used as a housekeeping gene, and the relative amount of mRNA was calculated by the 2^−ΔΔCT^ method.

### RNA sequencing

Total RNA was extracted from the ICC tissues using Qiagen kits. RNA sequencing was performed at the Molecular Biology Core Facilities, Dana Farber Cancer Institute (Boston, MA). The quality control of RNA-Seq raw data was performed by FastQC. After quality control, the low-quality bases and adaptors contamination was removed by Cutadapt. The quality of clean yield data was examined again by FastQC software. Next, the clean data were aligned to mouse reference genome mm10 by Hisat2 with a maximum of 2 mismatches per read. After data mapping, samtools and HTSeq-count were used to count the number of reads aligned to the gene features. The differentially expressed genes were identified by edgeR with a cutoff of |log(fold change)|>1 and p-value <0.01. Differentially expressed genes were annotated using Gene Ontology (GO), BIOCARTA, REACTOME, PID, and Kyoto Encyclopedia of Genes and Genomes (KEGG) public databases.

### Flow cytometry analysis

Cells were washed with the buffer, fixed, and permeabilized with FoxP3/Transcription Factor Staining Buffer Set (eBioscience/Thermo Fischer Scientific) to stain the intracellular markers. Harvested cells were incubated in Dulbecco’ s Modified Eagle Medium (DMEM) with a cell activation cocktail with brefeldin A (Biolegend) for 4 hours at 37°C. The cells were stained with the cell surface antibodies and intracellular marker in the buffer with brefeldin A. Anti-mouse CD16/32 antibody (clone 93, Biolegend, San Diego, California, USA) was added for FcR blockade and incubated for 5min at room temperature. After another washing step, antibodies for cell phenotyping were added, and cells were incubated for 40min at room temperature. The monoclonal antibodies used for flow cytometry analysis were CD8a (5H10-1), FoxP3 (FJK-16s), Cxcr3 (CXCR3-173), IFN-γ (XMG1.2), CD45 (30-F11), CD3e (145-2C11) and CD4 (RM4-5). The gating strategy of flow cytometry analysis is shown in **Fig. S11**.

### Imaging mass cytometry (IMC)

A tissue microarray (TMA) containing 63 cores, each 1.5 mm in diameter, was constructed; 54 of which were representative of the 18 mouse tumors based on the H&E review. Cores from a normal spleen were also included in the TMA as controls. To perform immunohistochemical staining with mass cytometry antibodies, the TMA slide was dewaxed in xylene and rehydrated in an alcohol gradient. Slides were incubated in Antigen Retrieval Agent pH 9 (Agilent^®^ S2367) at 96°C for 30 minutes; the slide was blocked with 3% BSA in PBS for 45 minutes at room temperature, followed by an overnight stain at 4°C with the antibody cocktail listed in **Table S3**. Cell-ID™ Intercalator-Ir (Standard Biotools PN 201192A) was used for DNA labeling. Ruthenium tetroxide 0.5% Aqueous Solution (Electron Microscopy Sciences PN 20700-05) was used as a counterstain. Images were acquired using a Hyperion Imaging System (Standard BioTools) at the Johns Hopkins Mass Cytometry Facility. Upon image acquisition, multi-layered ome.tiff images were generated for each core and exported using MCD™ Viewer (Standard BioTools). Using HALO 3.2, Area Quantification FL v2.1.2 was used to quantify the area density of individual IMC markers: CD8a, CD31, and Ir for DNA/Nuclear labeling. Area Quantification FL v2.1.2 was also employed to quantify the colocalized area density of IMC marker combinations CD3+TCF+, CD3+CD8+TCF+, and Ir for DNA/Nuclear labeling. All area density data were normalized by nuclear area density. The resulting data were analyzed by the Kruskall-Wallis test with pairwise comparisons using GraphPad Prism v9.2.0. Single-cell data obtained from segmented IMC images were analyzed following the workflow for IMC data (*47*), and plots were drawn in R-4.2.2.

### Statistical analysis

The χ2 (Chi-squared) or Fisher’s test was used to compare categorical variables, and the Mann-Whitney u test was utilized to compare two groups with quantitative variables. When the experimental cohort includes more than two groups, including quantitative variables, one-way ANOVA with Tukey’s multiple comparisons test was applied. The Kaplan-Meier method generated survival curves underlying the Log-Rank test and Cox proportional hazard model. The hazard ratio (HR) and 95% CI were calculated for statistical survival analyses for murine models. All analyses were performed using JMP Pro 11.2.0 (SAS Institute Inc., NC, USA), and data presented as mean ± S.E.M. Significant difference between experimental groups was determined when p-values were less than 0.05.

## Acknowledgments

The authors would like to express their sincere gratitude to Dr. M. Vader Heiden (MIT), Dr. S. Pillai (MGH), and Maksym Zarodniuk (University of Notre Dame) for valuable discussions, and Mark Duquette, Anna Khachatryan, and Sylvie Roberge for outstanding technical support.

## Author contributions

Conceptualization: JC, ZA, XL, SA, RKJ, DGD

Methodology: JC, ZA, XL, SA, AM, P-JL, MD, LLM, DF, PH, LS, NB, WJH

Investigation: JC, ZA, XL, SA, ZL, SS, AM, ZP, P-JL, MD, LZ, ZR, DS, KI

Visualization: JC, ZA, XL, P-JL, MD, LLM, DGD

Funding acquisition: DGD

Project administration: DGD

Supervision: DGD

Writing – original draft: JC, ZA, XL, DGD

Writing – review & editing: All authors

## Data and materials availability

All data, code, and materials used in the analysis are available electronically to any researcher. The imaging mass cytometry dataset has been deposited in Zenodo (DOI: 10.5281/zenodo.7439428). Reagents are available via materials transfer agreements (MTAs). All data are available in the main text or the supplementary materials.

## Supplementary Materials

Fig. S1. Standard chemotherapy converts ICB-resistant ICCs to ICB-responsive tumors, significantly delays tumor progression, and increases survival in mice.

Fig. S2. CTLA-4 blockade is critical for the efficacy of combined chemotherapy with ICB ICC by grafting 425 murine cells in C57Bl/6/FVB F1 mice.

Fig. S3. CTLA-4 blockade mediates the efficacy of GC/ICB therapy in ICC and increases CD8^+^Cxcr3^+^ CTL frequency in murine ICC.

Fig. S4. CD8^+^ T cells immunophenotyping in murine ICC.

Fig. S5. IMC analysis of ICC after GC-based therapies in orthotopic 425 murine ICC model.

Fig. S6. Bulk tissue RNA sequencing analysis of ICC after GC/dual ICB combination therapy in orthotopic 425 murine ICC model.

Fig. S7. Cxcr3 and its ligands are increased in ICC tissues after GC/ICB treatment in 425 murine ICC, and CXCR3 expression in human ICC, selective for T-cells, is a good prognostic factor.

Fig. S8. Experimental design for T-cell transfer experiment in mice bearing 425 murine ICC and treated with GC/ICB or GC alone.

Fig. S9. Cxcr3 in CD8 T-cells mediates the benefit of GC/ICB combination therapy in the 425 murine ICC model.

Fig. S10: Effect of ICB treatment scheduling on efficacy and toxicity.

Fig. S11: Gating strategies for flow cytometry.

Table S1. List of primary antibodies for flow cytometry and IF analyses.

Table S2. List of primers for qPCR analyses.

Table S3. List of antibodies used for IMC analyses.

Dataset S1. Bulk RNA sequencing of 425 murine ICC tissues after treatment.

