## Supplemental Figures for "Reprogramming Intrahepatic Cholangiocarcinoma Immune Microenvironment by Chemotherapy and CTLA-4 Blockade Enhances Anti-PD1 Therapy"

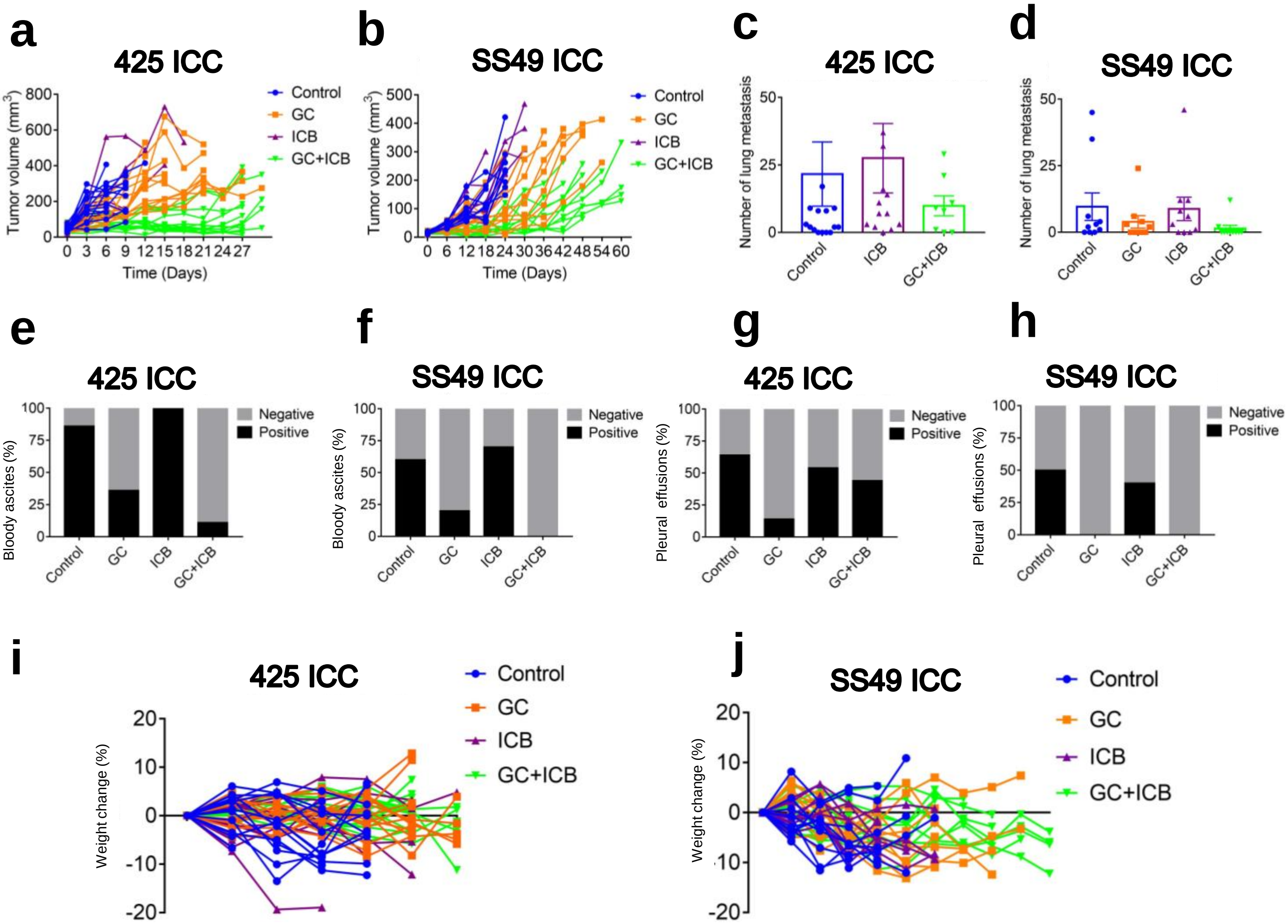


**Supplemental Figure S1: Standard chemotherapy converts ICB-resistant ICCs to ICB-responsive tumors, significantly delays tumor progression and increases survival in mice.** (**a**, **b**) Individual tumor curves for orthotopic ICCs after treatment: GC+ICB therapy induced a tumor growth delay that was significant superior to GC alone, while ICB alone was ineffective in 425 (**a**) and SS49 murine ICC models (**b**). (**c**, **d**) GC+ICB combined treatment significantly reduced ICC lung metastasis in 425 (**c**) and SS49 (**d**) murine ICC models. (**e-h**) GC+ICB combined treatment significantly reduced ICC-induced morbidity, by reducing ascites and pleural effusions in 425 (**e**, **g**) and SS49 (**f**, **h**) murine ICC models (**i**, **j**) Weight change after treatment with GC/ICB or GC or ICB alone versus control in 425 (**i**) and SS49 (**j**) murine ICC models. GC, gemcitabine plus cisplatin; ICB, immune checkpoint blockade (anti-PD-1 antibody plus anti-CTLA-4 antibody).

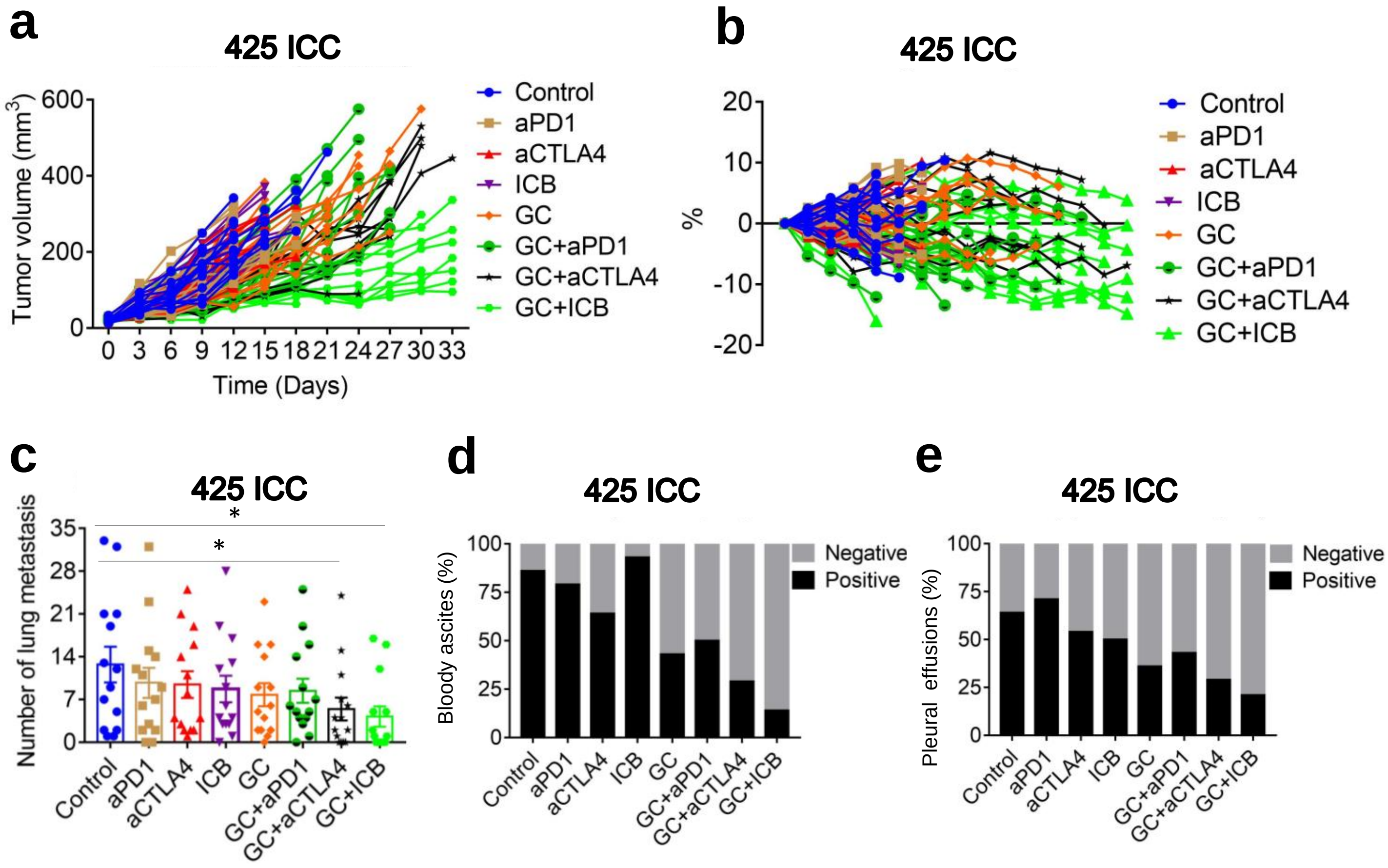


**Supplemental Figure S2: CTLA-4 blockade is critical for efficacy of combined chemotherapy with ICB in ICC by grafting 425 murine cells in C57Bl/6/129 F1 mice.** (**a**) Tumor curve of ICC after treatment: GC+ICB therapy induced a tumor growth delay that was significant superior to GC alone or even GC+aCTLA4, while ICB alone was ineffective in 425 and in SS49 ICC models. (**b**) Weight change demonstrated that GC+ICB is tolerance when compared with GC or ICB alone. (**c**) GC+ICB combined treatment significantly reduced ICC-induced morbidity, by reducing lung metastasis. (**d**) GC+ICB combined treatment significantly reduced ICC-induced morbidity, by reducing ascites and (**e**) pleural effusions. GC: gemcitabine plus cisplatin; ICB: anti-PD-1 antibody plus anti-CTLA-4 antibody. *$p < 0.05$ from Unpaired t test.

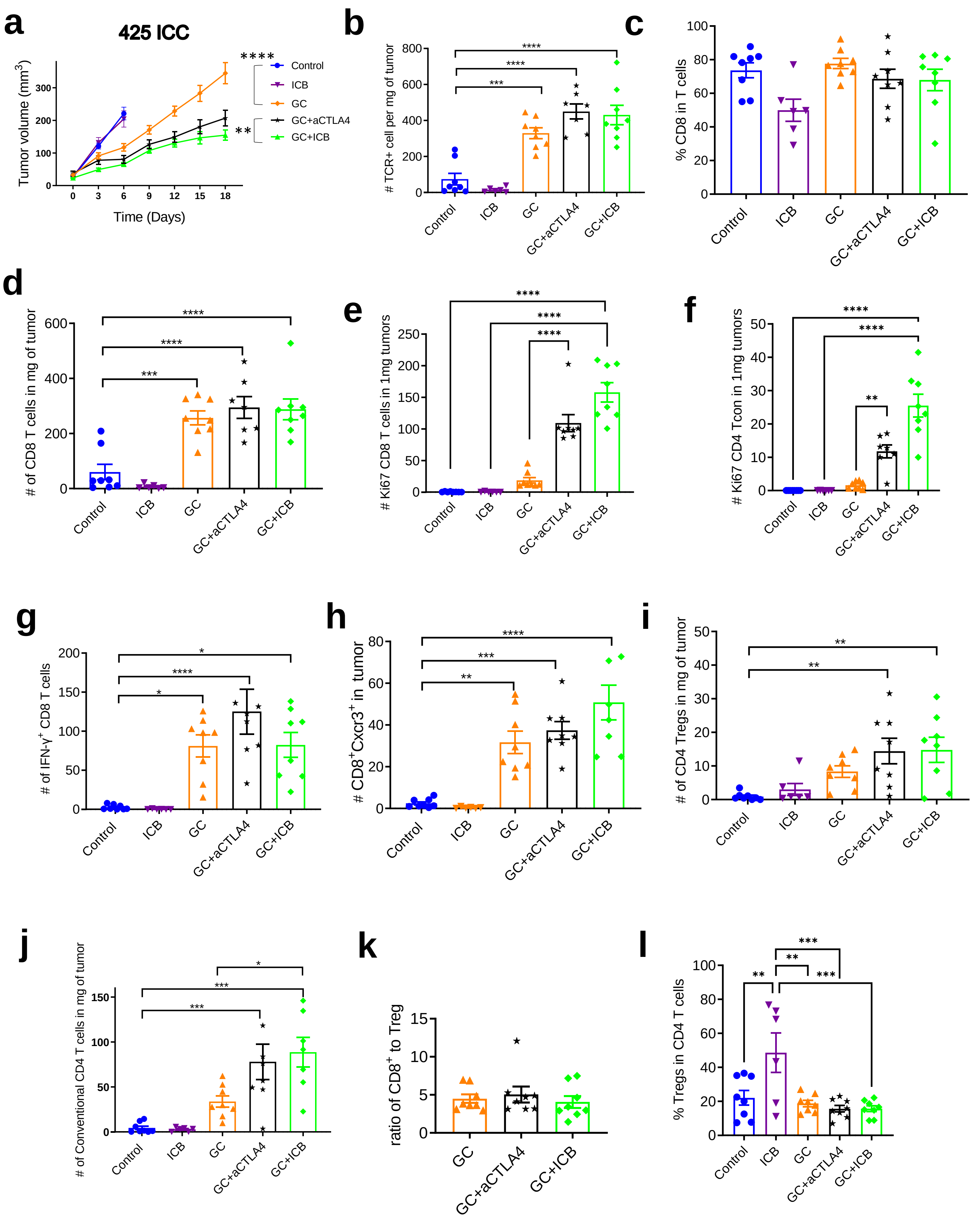

a
425 ICC
Tumor volume (mm3)
Time (Days)
Control
ICB
GC
GC+aCTLA4
GC+ICB
b
### TCR+ cell per mg of tumor
c
% CD8 in T cells
d
### of CD8 T cells in mg of tumor
e
### Ki67 CD8 T cells in 1mg tumors
f
### Ki67 CD4 Tcon in 1mg tumors
g
### of IFN-γ+ CD8 T cells
h
### CD8+Cxcr3+ in tumor
i
### of CD4 Tregs in mg of tumor
j
### of Conventional CD4 T cells in mg of tumor
k
ratio of CD8+ to Treg
l
% Tregs in CD4 T cells

**Supplemental Figure S3: CTLA-4 blockade mediates the efficacy of GC/ICB therapy in ICC and increases $CD8^+Cxcr3^+$ CTL frequency in murine ICC.** (**a**) Tumor growth kinetics after treatment in orthotopic 425 murine ICC model: GC/anti-CTLA4 Ab combination is significantly superior to GC alone and in delaying tumor growth; addition of anti-PD1 Ab to GC/anti-CTLA4 Ab (GC/ICB) induces a greater delay in tumor growth. (**b-l**) Immunophenotyping of treated ICC tissues at days 10 (control and ICB groups) and 20 (GC-containing groups). $TCR^+$ TIL numbers were higher in all GC-treated groups (**b**). $CD8^+$ T cells represented approximatively 60% of TILs in all groups (**c**), and their numbers (**d**) were increased in all GC-treated groups. The numbers of $Ki67^+CD8^+$ (**e**) and $Ki67^+CD4^+$ (**f**) were increased in GC/anti-CTLA4 and GC/dual ICB groups. The numbers of $CD8^+IFN\text{-}\gamma^+$ (**g**) and $CD8^+Cxcr3^+$ (**h**) were increased in all GC-treated groups. Numbers of $CD4^+FoxP3^+$ Treg (**i**) and $CD4^+$ conventional T cells (**j**) proportionally increased in all GC-treated groups. Ratio of CD8 to Treg were no significant difference between GC-treated groups (**k**). The frequency of Tregs in all $CD4^+$ T cells in GC-treated groups were not increased (**l**). *$p < 0.05$; **$p < 0.01$; ***$p < 0.001$ ; ****$p < 0.0001$ from Dunnett's multiple comparisons test (**a**) and from Tukey's multiple comparisons test (**b-l**). GC: gemcitabine plus cisplatin; ICB: anti-PD-1 antibody plus anti-CTLA-4 antibody; ICC, intrahepatic cholangiocarcinoma; CTL, cytotoxic T lymphocyte; Treg, regulatory T cell; TCR, T cell receptor; TIL, tumor-infiltrating lymphocyte (TIL).

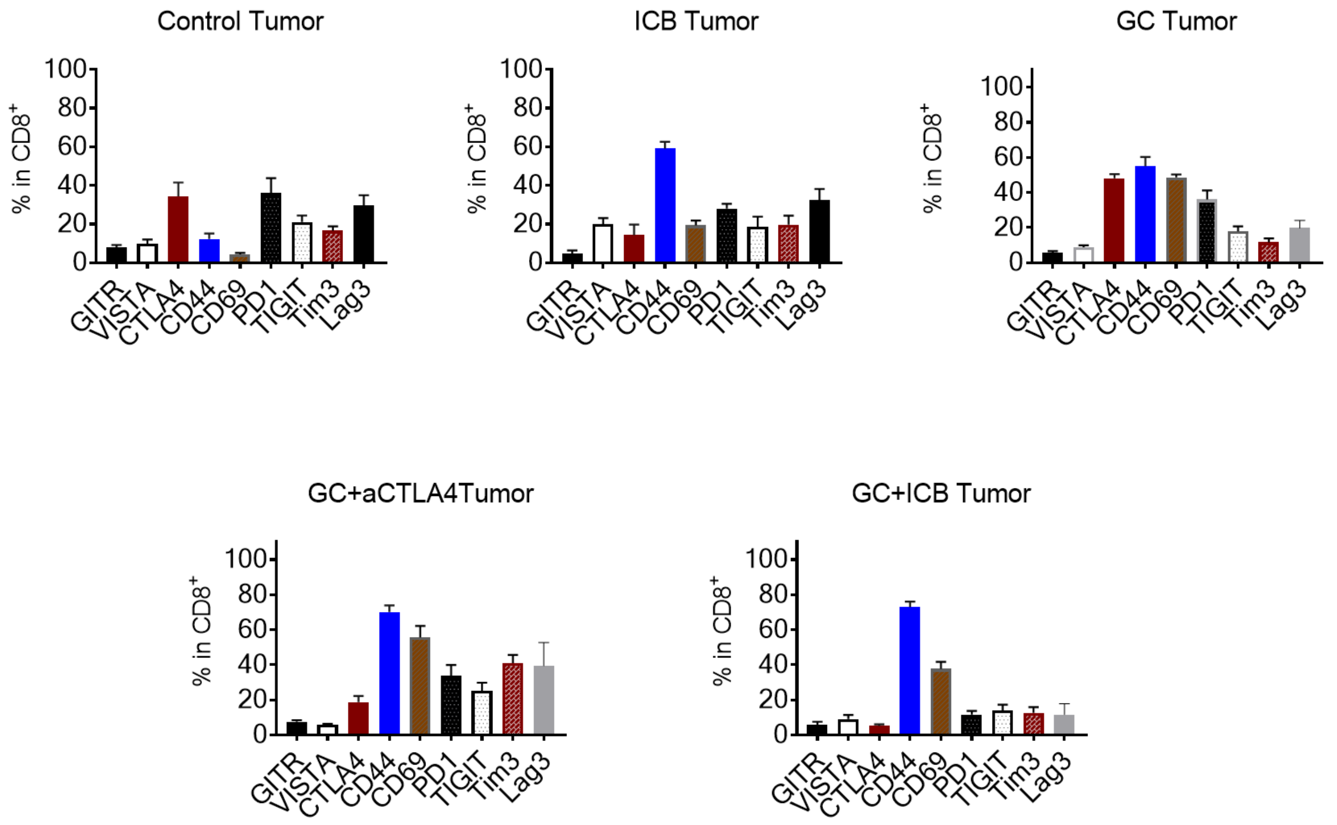


**Supplemental Figure S4: CD8+ T cells immunophenotyping in murine ICC.** Immunophenotyping of CD8+ T cells from treated ICC tissues with immune checkpoint molecules and activation markers. GC: gemcitabine plus cisplatin; ICB: anti-PD-1 antibody plus anti-CTLA-4 antibody; ICC, intrahepatic cholangiocarcinoma.

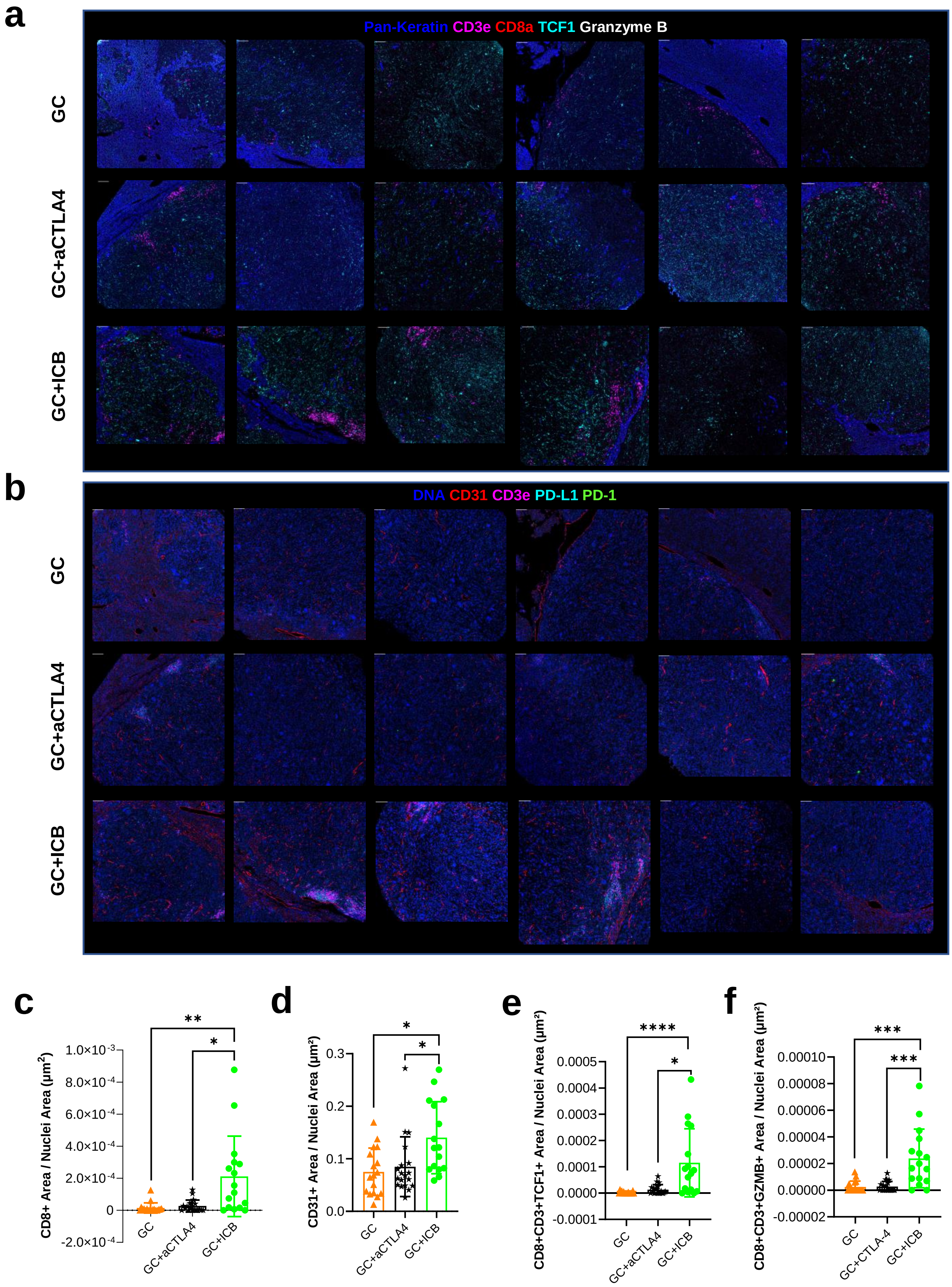
a
Pan-Keratin CD3e CD8a TCF1 Granzyme B
GC
GC+aCTLA4
GC+ICB
b
DNA CD31 CD3e PD-L1 PD-1
GC
GC+aCTLA4
GC+ICB
c
CD8+ Area / Nuclei Area (µm²)
1.0×10-3
8.0×10-4
6.0×10-4
4.0×10-4
2.0×10-4
0
-2.0×10-4
**
*
GC
GC+aCTLA4
GC+ICB
d
CD31+ Area / Nuclei Area (µm²)
0.3
0.2
0.1
0.0
*
*
GC
GC+aCTLA4
GC+ICB
e
CD8+CD3+TCF1+ Area / Nuclei Area (µm²)
0.0005
0.0004
0.0003
0.0002
0.0001
0.0000
-0.0001
****
*
GC
GC+aCTLA4
GC+ICB
f
CD8+CD3+GZMB+ Area / Nuclei Area (µm²)
0.00010
0.00008
0.00006
0.00004
0.00002
0.00000
-0.00002
***
***
GC
GC+CTLA-4
GC+ICB

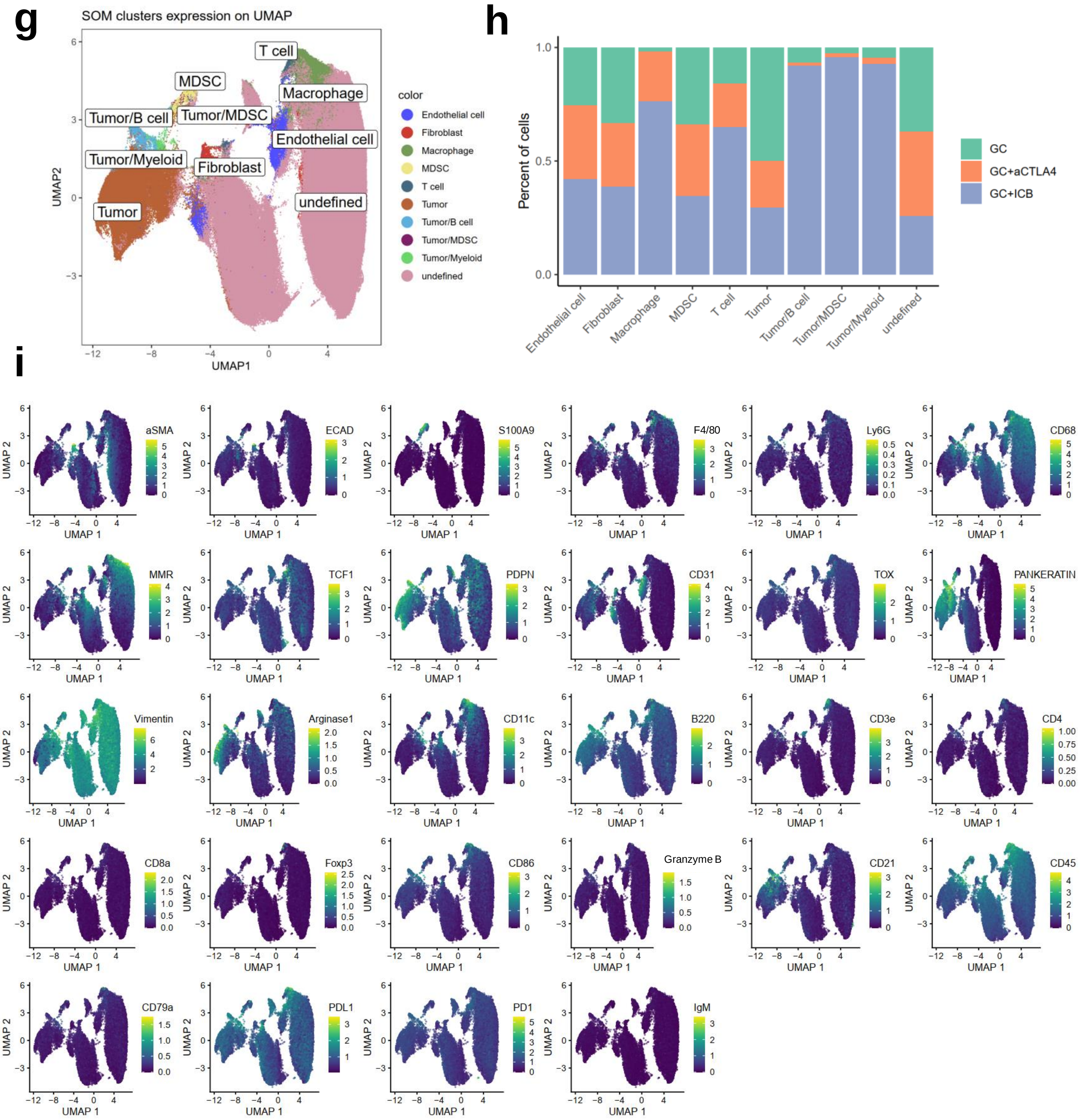


**Supplemental Figure S5: IMC analysis of ICC after GC-based therapies in orthotopic 425 murine ICC model.** (**a-b**) Representative multicolored images for every sample in the tissue microarray for Pan-Keratin, CD3e, CD8a, TCF1, and Granzyme B (**a**), DNA, CD31, CD3e, PD-L1, and PD-1 (**b**), N=6 mice/group. (**c,d**) GC combined with ICB can increase CD8+ and CD31+ cells in ICC tissues quantified by IMC. (**e,f**) The abundance of CD8+CD3+TCF1+ cells and CD8+CD3+GZMB+ cells assessed by IMC showed a significant increase in GC/ICB combination group. (**g**) UMAP plot shows the self-organizing map clusters with annotations based on cell segmentation data from IMC images. Tumor/B cell, Tumor/MDSC, Tumor/Myeloid mean the tumor next to B cell, tumor next to MDSC, and tumor next to myeloid cells. (**h**) The proportion of each cell cluster among all treatment groups. (**i**) UMAP plots showing the expression of markers included in the IMC panel. $*p < 0.05$; $**p < 0.01$; $***p < 0.001$; $****p < 0.0001$, Kruskal-Wallis Test and 3 high-power fields for each sample. GC: gemcitabine plus cisplatin; ICB: anti-PD-1 antibody plus anti-CTLA-4 antibody.

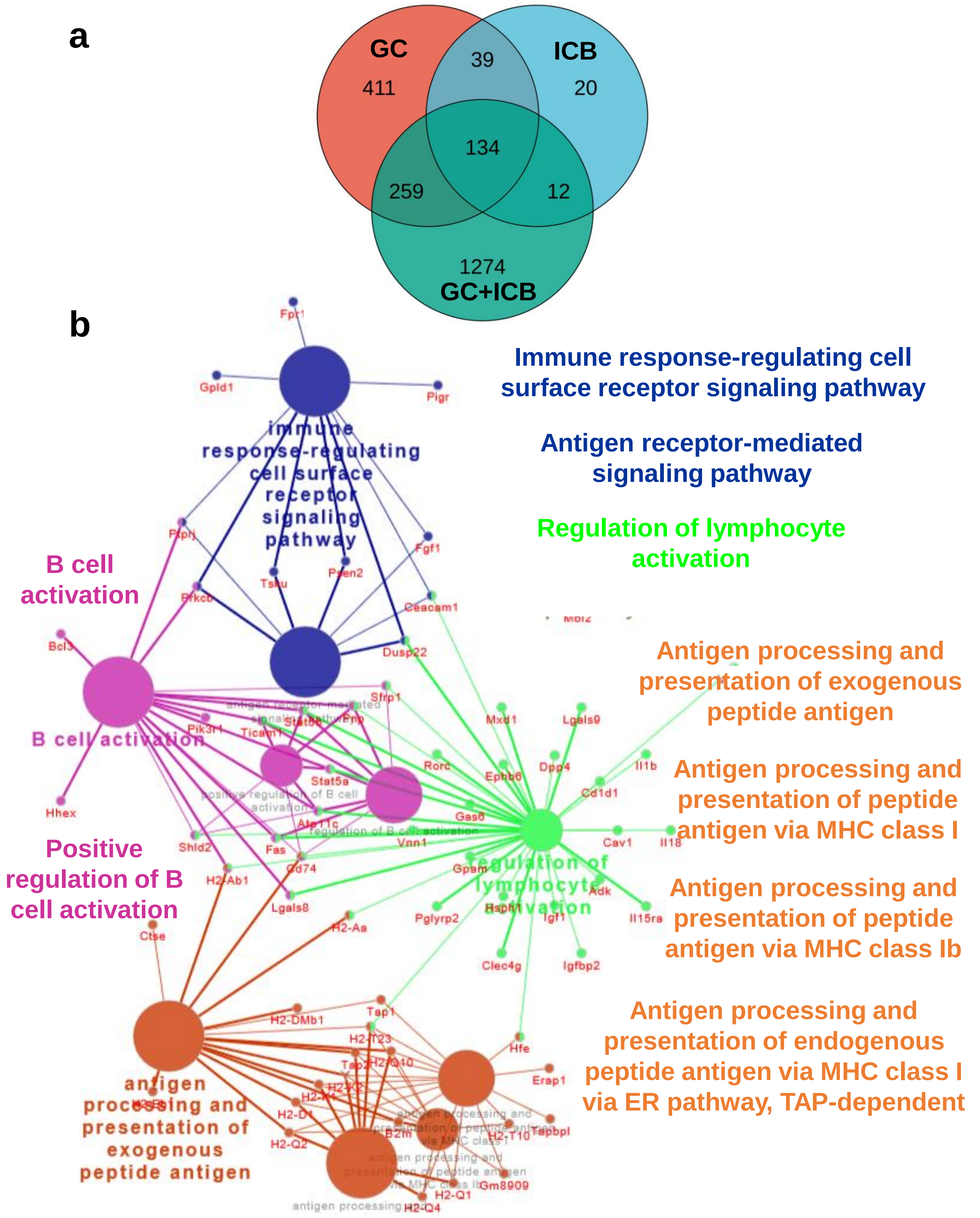


**Supplemental Figure S6: Bulk tissue RNA sequencing analysis of ICC after GC/dual ICB combination therapy in orthotopic 425 murine ICC model.** (**a**) Differentially expressed genes between groups. (**b**) Inflammatory pathways activated in ICC tissue after GC/dual ICB versus IgG control treatment (KEGG database geneset enrichment analysis).

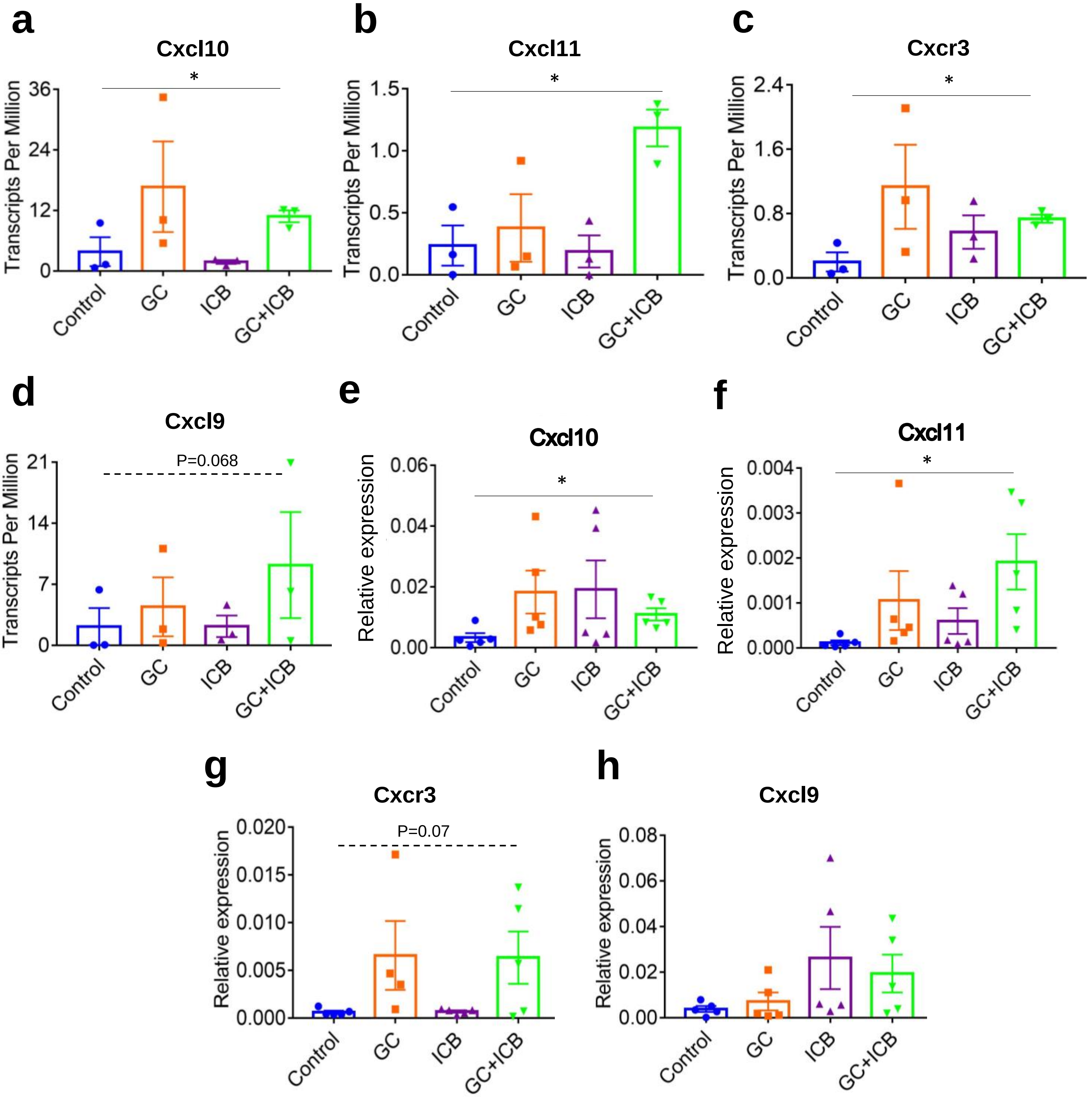


**Supplemental Figure S7: Expression levels of Cxcr3 and its ligands are increased in ICC tissues after GC/dual ICB treatment in 425 murine ICC, and CXCR3 expression in human ICC, selective for T cells, is a good prognostic factor.** (**a-d**) RNAseq analysis showing a significant increase in the expression of Cxcl10 (**a**), Cxcl11 (**b**) and Cxcr3 (**c**), and a trend for increased Cxcl9 expression (**d**) after GC/ICB versus control (n=3). (**e,f**) Quantitative PCR analysis for Cxcl10 (**e**) and Cxcl11 (**f**) showing a significant increase in gene expression in GC/ICB versus control group (n=6). (**g,h**) Quantitative PCR analysis for Cxcr3 (**g**) and Cxcl9 (**h**) showing a trend for increased Cxcr3 gene expression (n=6). *p<0.05 from Unpaired t test (**a-c, e,f**).

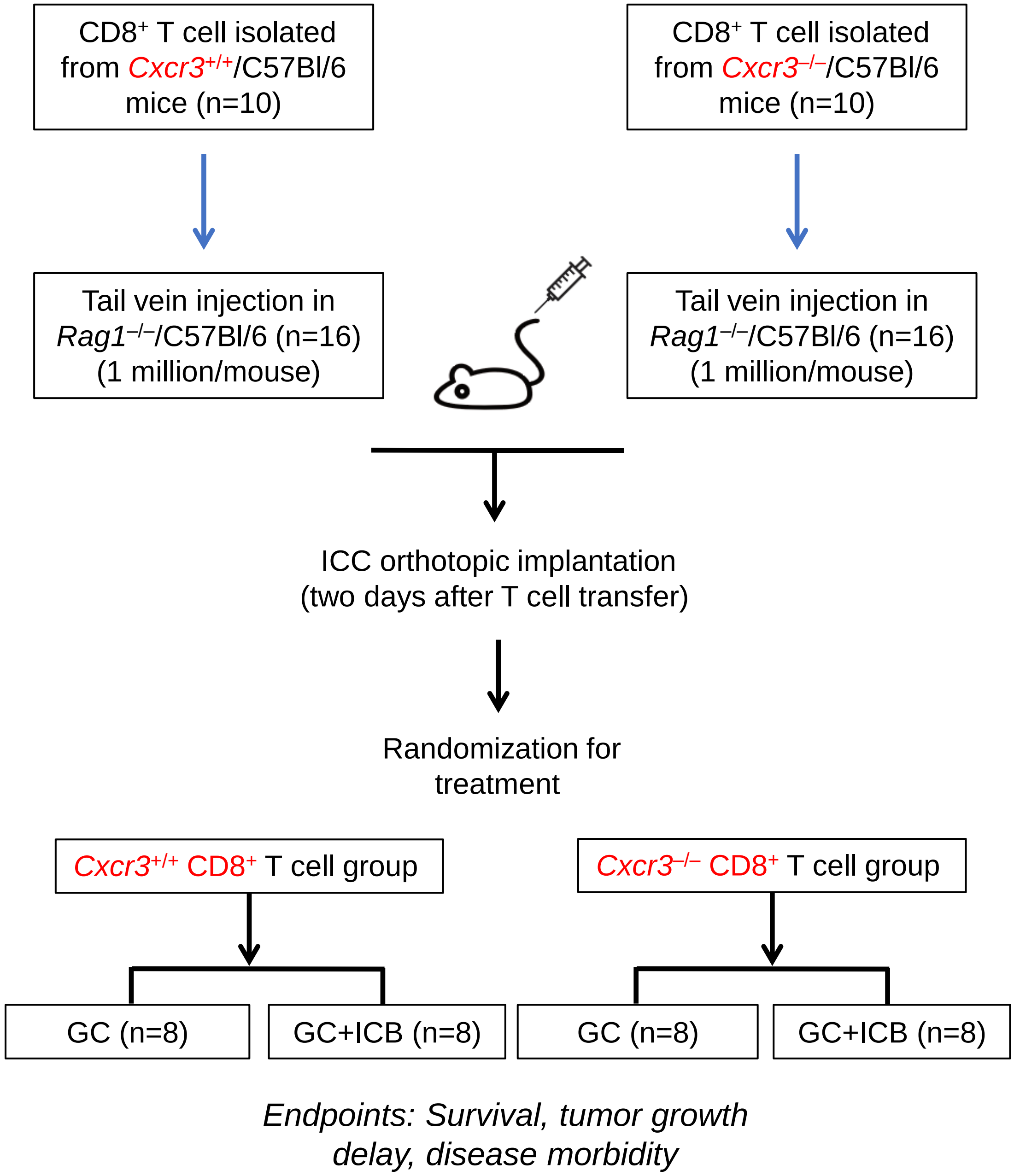


**Supplemental Figure S8: Experimental design for T cell transfer experiment in mice bearing 425 murine ICC and treated with GC/ICB or GC alone.** Mice with established orthotopic 425 tumor were treated with GC (twice/week) alone or GC (twice/week) + ICB (every 3 days).

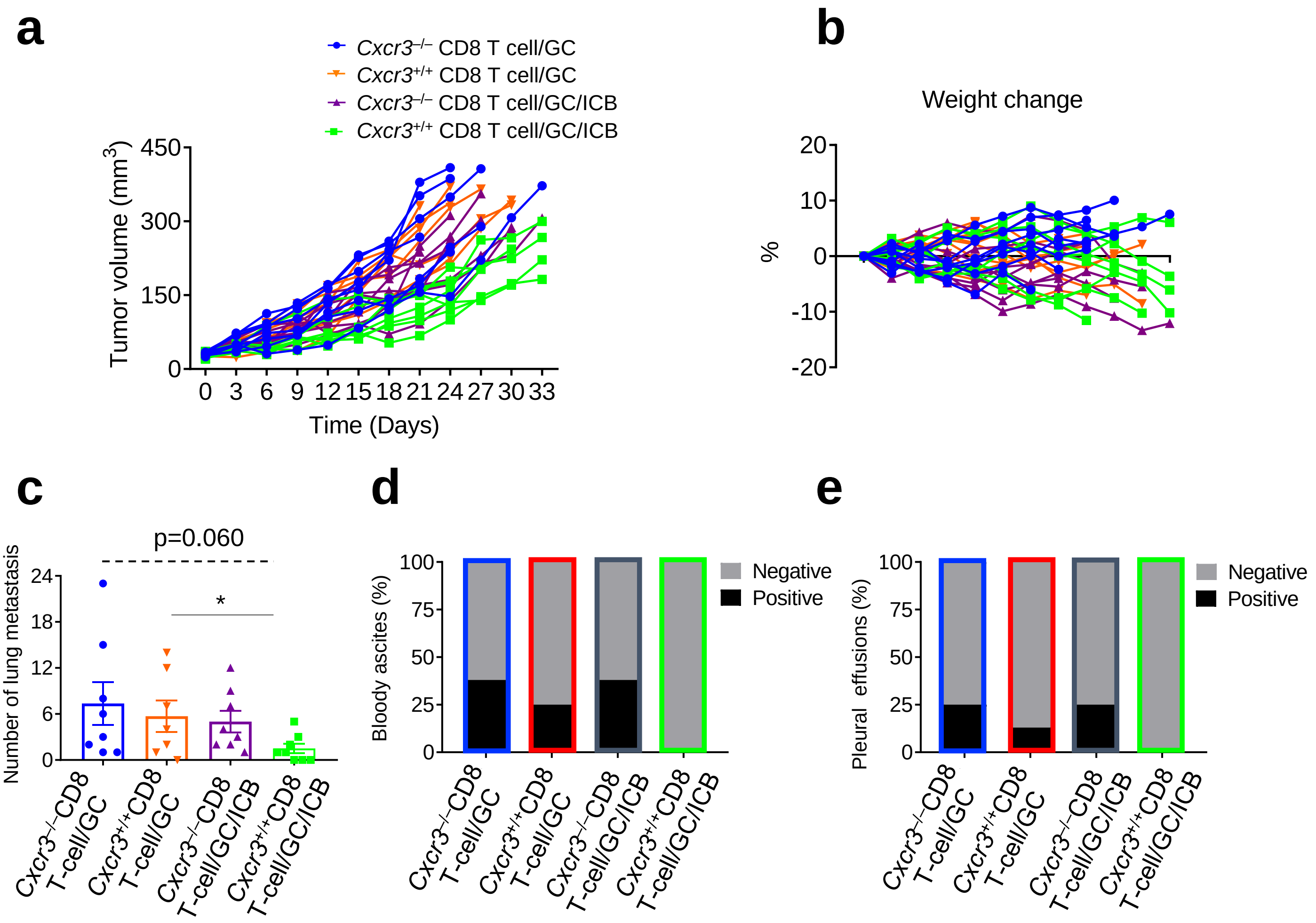


**Supplemental Figure S9: CXCR3 in CD8 T cells mediates the benefit of GC/ICB combination therapy in orthotopic 425 murine ICC model.** (**a-e**) Outcomes of GC+ICB versus GC alone therapy in 425 ICC-bearing *Rag1*$^{-/-}$/C57Bl/6 mice which received CD8$^{+}$ T cell transfer from *Cxcr3*$^{-/-}$/C57Bl/6 or *Cxcr3*$^{+/+}$/C57Bl/6 mice (n=8 mice). Individual tumor growth kinetics (**a**), changes in body weight (**b**), lung metastases (**c**), bloody ascites (**d**), and pleural effusions (**e**). GC: gemcitabine plus cisplatin; ICB: anti-PD-1 antibody plus anti-CTLA-4 antibody. *$p < 0.05$ from Unpaired t test (**c**).

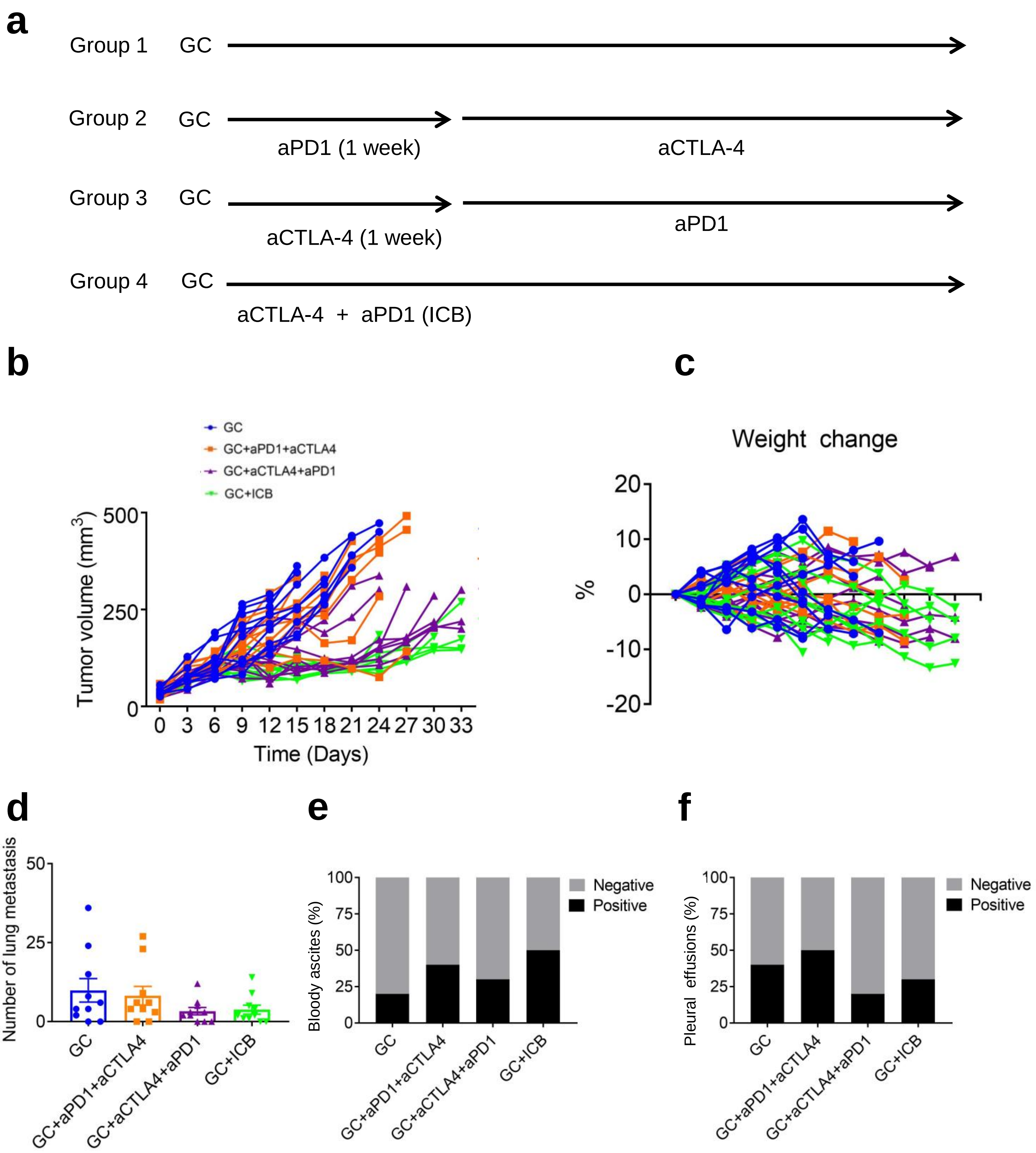


**Supplemental Figure S10: Effect of ICB treatment scheduling on efficacy and toxicity.** (**a**) Treatment groups and sequencing of treatments; GC, gemcitabine/cisplatin. (**b**) Individual tumor growth kinetics. (**c**) Changes in body weight. (**d**) Lung metastatic burden. (**e**, **f**) Incidence of bloody ascites (**e**), and pleural effusions (**f**).

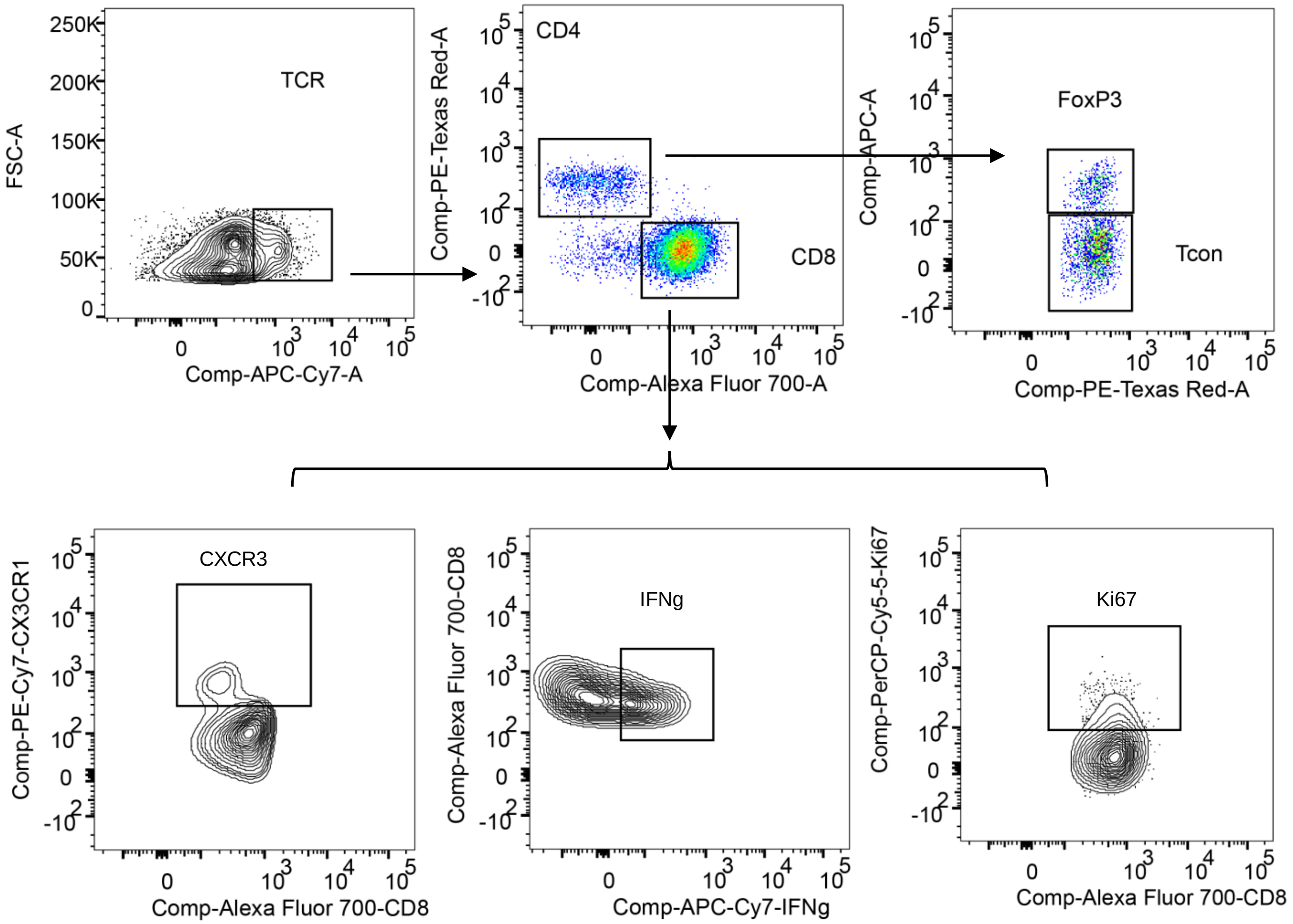


**Supplemental Figure S11: Gating strategies for flow cytometry.**
