## Supplementary material for "Reprogramming Intrahepatic Cholangiocarcinoma Immune Microenvironment by Chemotherapy and CTLA-4 Blockade Enhances Anti-PD1 Therapy": Table S1

| **Table S1: Primary antibodies** | |
| --- | --- |
| **Name** | **Source and catalog number** |
| TCR-BV711 | BioLegend Cat. No. 109243 |
| CD39-PECY7 | BioLegend Cat. No. 143806 |
| Ki67-Percyp | BioLegend Cat. No. 652424 |
| CD11b-bv785 | BioLegend Cat. No. 101243 |
| CD206-BV420 | BioLegend Cat. No. 141717 |
| BV421-FoxP3 | BioLegend Cat. No. 126419 |
| IFNG-Rb-APC | [BioLegend Cat. No. 113606](https://www.biolegend.com/en-us/products/apc-anti-mouse-ifn-y-r-b-chain-antibody-16014) |
| CD119 | [BD Biosciences, Cat. No. 2740545](https://www.bdbiosciences.com/us/reagents/research/antibodies-buffers/immunology-reagents/anti-human-antibodies/cell-surface-antigens/bv786-rat-anti-mouse-cd119-gr20/p/740897) |
| BV510-IFNG | BioLegend Cat. No. 505842 |
| FiTC-anti-mouse IL2 | [BioLegend Cat. No. 503806](https://www.biolegend.com/en-us/products/fitc-anti-mouse-il-2-antibody-952) |
| Mouse CD31 | Millipore Cat. No. MAB1398Z |
| Mouse α-SMA | SIGMA Cat. No. C6198 |
| Mouse CD8 | Biorbyt Cat. No. orb348907 |
