## Supplementary material for "Reprogramming Intrahepatic Cholangiocarcinoma Immune Microenvironment by Chemotherapy and CTLA-4 Blockade Enhances Anti-PD1 Therapy": Table S2

| **Table S2: Primers for PCR** | |
| --- | --- |
| **Primers** | **Sequence** |
| muActin F | GGCTGTATTCCCCTCCATCG |
| muActin R | CCAGTTGGTAACAATGCCATGT |
| 126mGAPDH F | AGGTCGGTGTGAACGGATTTG |
| 126mGAPDH R | TGTAGACCATGTAGTTGAGGTCA |
| muCXCL11 F | AGCTGCTCAAGGCTTCCTTA |
| muCXCL11 R | AGTAACAATCACTTCAACTTTGTCG |
| muCXCL9 F | GTTCGAGGAACCCTAGTGATAAGG |
| muCXCL9 R | CCTCGGCTGGTGCTGATG |
| muCXCL10 F | CCAAGTGCTGCCGTCATTTTC |
| muCXCL10 F | GGCTCGCAGGGATGATTTCAA |
| muCXCR3 F | TTGCCCTCCCAGATTTCATC |
| muCXCR3 R | TGGCATTGAGGCGCTGAT |
