## Supplementary material for "Reprogramming Intrahepatic Cholangiocarcinoma Immune Microenvironment by Chemotherapy and CTLA-4 Blockade Enhances Anti-PD1 Therapy": Table S3

| **Table S3: IMC panel** |
| --- |

| **Mass** | **Metal** | **Antigen** | **Clone** | **Conc. (μg/mL)** | **Source** | **Custom** |
| --- | --- | --- | --- | --- | --- | --- |
| 96-104 | Ru | Counterstain |  | 20 | Electron Microscopy Sciences |  |
| 113 | In | αSMA | 1A4 | 20 | Cell Signaling Technology® | X |
| 115 | In | E-cadherin | 24E10 | 20 | Cell Signaling Technology® | X |
| 141 | Pr | S100A9 | D3U8M | 20 | Cell Signaling Technology® | X |
| 142 | Nd | F4/80 | D2S9R | 10 | Cell Signaling Technology® | X |
| 143 | Nd | Ly6G | 1A8 | 20 | Biolegend® | X |
| 144 | Nd | CD68 | E3O7V | 20 | Cell Signaling Technology® | X |
| 146 | Nd | CD206 | E6T5J | 13.33 | Cell Signaling Technology® | X |
| 149 | Sm | TCF1/TCF7 | C63D9 | 20 | Cell Signaling Technology® | X |
| 150 | Nd | Podoplanin | 8.1.1 | 20 | Biolegend® | X |
| 152 | Sm | CD31 | D8V9E | 20 | Cell Signaling Technology® | X |
| 153 | Eu | Tox/Tox2 | E6I3Q | 20 | Cell Signaling Technology® | X |
| 154 | Eu | Pan-Keratin | C11 | 20 | Cell Signaling Technology® | X |
| 155 | Gd | Vimentin | D21H3 | 20 | Cell Signaling Technology® | X |
| 156 | Gd | Arginase-1 | D4E3M | 13.33 | Cell Signaling Technology® | X |
| 158 | Gd | CD11c | D1V9Y | 20 | Cell Signaling Technology® | X |
| 160 | Gd | B220 | RA3-6B2 | 20 | Cell Signaling Technology® | X |
| 161 | Dy | CD3e | E4T1B | 20 | Cell Signaling Technology® | X |
| 162 | Dy | CD4 | EPR19514 | 20 | Abcam | X |
| 163 | Dy | CD8a | D4W2Z | 20 | Cell Signaling Technology® | X |
| 165 | Ho | Foxp3 | D6O8R | 20 | Cell Signaling Technology® | X |
| 166 | Er | CD86 | E5W6H | 8 | Cell Signaling Technology® | X |
| 167 | Er | Granzyme B | E5V2L | 20 | Cell Signaling Technology® | X |
| 170 | Er | CD21 | SP186 | 20 | Abcam | X |
| 172 | Yb | CD45 | D3F8Q | 20 | Cell Signaling Technology® | X |
| 173 | Yb | CD79a | EPR26537-114 | 20 | Abcam | X |
| 174 | Yb | PD-L1 | D5V3B | 20 | Cell Signaling Technology® | X |
| 175 | Lu | PD-1 | EPR20665 | 20 | Abcam | X |
| 176 | Yb | IgM | II/41 | 20 | eBioscience™ | X |
| 191 | Ir | DNA 1 |  |  | Standard BioTools™ |  |
| 193 | Ir | DNA 2 |  |  | Standard BioTools™ |  |
| 195 | Pt | Plasma Membrane 2 |  |  | Standard BioTools™ |  |
| 196 | Pt | Plasma Membrane 3 |  |  | Standard BioTools™ |  |
| 198 | Pt | Plasma Membrane 4 |  |  | Standard BioTools™ |  |
